# Antiviral viral compound from *Streptomyces ghanaensis* like strain against white spot syndrome virus (WSSV) of shrimp

**DOI:** 10.1101/340265

**Authors:** T. Rajkumar, M. Manimaran, G. Taju, S. Vimal, S. Abdul Majeed, K. Kannabiran, S. Sivakumar, K.M. Kumar, S. Madhan, A.S. Sahul Hameed

**Author notes:** **Corresponding author** Dr. A.S. Sahul Hameed.

## Abstract

Actinomycetes isolates collected from different environments were screened for antiviral activity against WSSV. One isolate designated as CAHSH-2 showed antiviral activity against WSSV at the concentration of 0.2 mg per shrimp. The laboratory trial of determining antiviral activity of ethyl acetate extract (EtOAcE) of CAHSH-2 against WSSV was carried out 21 times since 2014. CAHSH-2 isolate which showed antiviral activity was characterized and identified as *Streptomyces ghanaensis* like strain. Among the five fractions obtained from EtOAcE of potential actinomycetes isolate, F1 was found to have strong antiviral activity. The F1A and F1B sub-fractions from F1 fraction were subjected to GC-MS, FTIR, ^1^H and ^13^C NMR analyses and, the compounds identified were di-n-octyl phthalate and bis (2-methylheptyl) phthalate, respectively. Among these compounds, di-n-octyl phthalate showed strong antiviral activity against WSSV. Molecular docking studies revealed that di-n-octyl phthalate was found to have high binding affinity with VP26 and VP28 proteins of WSSV, whereas the bis (2-methylheptyl) phthalate showed low binding affinity with VP26 and VP28. The antiviral activity of EtOAcE of actinomycetes against WSSV was confirmed by PCR, RT-PCR, Western blot and ELISA. The EA extract of active isolate was found to be non-toxic to *Artemia*, post-larvae and adult *Litopenaeus vannamei*.

**Importance:** White spot syndrome virus (WSSV) is an important shrimp viral pathogen and responsible for huge economic loss to shrimp culture industry worldwide including India. The global loss due to WSSV has been estimated about USD 10 billion and the loss continues at the same extent even now. Various strategies have been followed to prevent or control diseases of aquatic animals. In spite of various preventive and control strategies, WSSV has been still persisting for more than two decades. No control strategies have so far been evolved to put a break to WSSV. In this situation, an attempt was made in the present work to screen some actinomycetes isolates for antiviral activity against WSSV. Among these isolates, one isolate identified as *Streptomyces ghanaensis* like isolate CAHSH-2 showed activity against WSSV. This article gives the information about the antiviral compound against WSSV and the mechanism of viral inhibition.

## 1. Introduction

Disease is now considered to be a major limiting factor in aquaculture production worldwide. The economic loss due to diseases has been estimated about US$ 1 billion in developing countries of Asia in 1990 alone. Since then, losses have increased. A 1995 estimate suggests that aquatic animal disease and environment-related problems may cause annual losses to aquaculture production in Asian countries to the tune of more than US$ 2 billion per year. According to recent reports, global losses due to shrimp disease are more than US$ 2.8 billion. In India, the loss due to white spot syndrome virus (WSSV) has been estimated about USD 150 million per year, and the loss continues even now to the same extent (1). Viral and bacterial diseases are very important since they are responsible for huge economic loss in aquaculture systems worldwide. Although information on biology and culture aspects of Indian cultivable species of fish, shrimp and prawn is available, studies on diseases of aquatic animals in Indian aquaculture system and their control and prevention are very few. The important viral and bacterial pathogens reported in Indian aquaculture system are WSSV, infectious hypodermal and hematopoietic necrosis (IHHNV), monodon baculovirus (MBV), hepatopancreatic necrosis virus (HPV), fish nodavirus, *Macrobrachium rosenbergii* nodavirus (MrNV), extra small virus, *Vibrio harveyi, V. anguillarum, Aeromonas hydrophila, A. caviae* and *Edwardsiella tarda*. A surveillance program supported by National Fisheries Development Board of ICAR, India is being carried out to monitor aquaculture systems throughout the country for emerging diseases in shrimp, prawn and fish. As a result of this program which is now into its third year, the occurrence of *Enterocytozoon hepatopenaei* (2, 3) and infectious myonecrosis virus (IMNV) in pond-reared *L. vannamei* (4), cyprinid herpesvirus-2 in goldfish (5) and tilapia lake virus in Nile tilapia (6) has been reported. In the field condition, various control strategies such as improvement of environmental conditions, stocking of specific pathogen-free shrimp post-larvae and augmentation of disease resistance by oral immunostimulants, are being applied to control WSSV infection. But none of these methods provided effective protection to shrimp against WSSV infection.

Actinomycetes are potential microorganisms that are capable of producing a variety of antiviral compounds working against viral pathogens of human, higher animals, aquatic animals and plants. (7) have isolated bioactive compounds from actinomycetes possessing high antiviral activity against acyclovir-resistant herpes simplex virus at non-cytotoxic concentration. Promising non-plant derived glycan-targeting compounds such as the mannose-specific pradimicin-A (PRMA) extracted from the actinomycete strain *Actinomadura hibisca*, showed strong binding capacity with human immunodeficiency virus (HIV) (8). Actinohivin (AH) is a potent anti-HIV lectin that is produced by an actinomycete, *Longispora albida* gen. nov, sp. nov (9, 10). (11) reported that Benzastatin C, a 3-chloro-tetrahydroquinolone alkaloid from *Streptomyces nitrosporeus*, showed antiviral activity in a dose-dependant manner against herpes simplex virus type 1 (HSV-1), herpes simplex virus type 2 (HSV-2), and vesicular stomatitis virus (VSV). (12) have isolated a novel anti-influenza virus compound (JBIR-68) from *Streptomyces* sp. RI18 and this compound inhibited influenza virus growth in plaque assays. A bioactive compound, xiamycin D isolated from *Streptomyces* sp (#HK18) culture inhabiting the topsoil in a Korean solar saltern was tested for antiviral activity against porcine epidemic diarrhea virus (PEDV) and results showed that xiamycin D had the strongest inhibitory effect on PEDV replication (13). Methylelaiophylin, an antiviral compound isolated from *Streptomyces melanosporofaciens* showed antiviral activity against Newcastle disease virus (NDV) (14).

Work on application of actinomycetes for aquatic animal health is very limited in India. (15) have isolated twenty-five isolates of actinomycetes and tested them for their ability to reduce WSSV-infection in shrimp. The antiviral activity of furan-2-yl acetate (C6H6O3) extracted from Streptomyces VITSDK1 spp. was studied in cultured Sahul Indian Grouper Eye (SIGE) cells infected with fish nodavirus (FNV). This compound (20 μg mL^−1^) effectively inhibited the replication of the FNV, and the viral titre was reduced from 4.3 to 2.45 log TCID50 mL^−1^ on treatment (16). (17) reported the antiviral activity of crude extract of marine *Streptomyces carpaticus* MK-01 isolated from seawater against viral hemorrhagic septicemia virus of fish. Based on the above studies, an attempt was made in the present study to screen a group of actinomycetes collected from different habitats for antiviral activity against WSSV of shrimp.

## 2. Materials and methods

### 2.1. Collection and maintenance of experimental animals

Marine shrimp, *Litopenaeus vannamei* (10–15 g body weight) were collected from grow-out ponds located at Nellore, AP or Ponneri, TN and maintained in 1000 l aquarium tanks containing seawater with salinity between 15 and 20 ppt (50 shrimp per aquarium tank) at room temperature which ranged from 27 to 30°C for 5 days prior to experiments. The shrimp were fed with commercial pellet feed. Seawater was obtained from the Research Centre of Central Institute of Brackishwater Aquaculture, Muttukadu near Chennai. Seawater was pumped from the adjacent sea and allowed to sediment, thus removing sand and other particulate matter before use for shrimp. Five shrimps were randomly selected and screened for WSSV by PCR with the primers designed by Yoganandhan *et al*. (18).

### 2.2 White spot syndrome virus (WSSV)

The WSSV suspension was prepared from WSSV-infected shrimp *L. vannamei* with clinical signs of lethargy, reddish colouration and white spots. The infected shrimp were collected from shrimp farms located near Nellore, India. Hemolymph was drawn directly from the heart of infected shrimp using sterile syringes. The hemolymph was centrifuged at 3000 *g* for 20 min at 4°C and the supernatant was collected. Then the collected supernatant was recentrifuged at 8000 *g* for 30 min at 4°C. The final supernatant fluid was then filtered through a 0.4 μm filter before storage at −20°C, until used in infectivity experiments. Prior to storage, the presence of WSSV was confirmed by PCR using primers designed by Yoganandhan *et al*. (18). Bioassay experiment was carried out to check the virulence of WSSV by injecting the viral suspension intramuscularly in healthy shrimp. This viral suspension was used for screening actinomycetes for antiviral activity against WSSV.

### 2.3. Collection of Actinomycetes isolates

In the present study, actinomycetes isolates were isolated from different environments like marine sediment, plants, effluents from industrial areas and freshwater samples using standard isolating procedures and recommended culture media (19, 20, 21). *Streptomyces ghanaensis* and *S. viridosporus* used as reference strains were obtained from IMTECH Culture Collection, Chandigarh, India and used for comparison to characterize the isolates of actinomycetes.

### 2.4. Extraction of metabolites

Actinomycetes isolates were grown separately in different media such as starch casein broth (SCB) and production medium-I (PM-I), PM-II, and PM-III to select the most suitable medium. The cultures were incubated at 28°C in a rotary shaker. The cultures were harvested after 7 days and filtered through Whatman No.1 filter paper. The culture filtrates were centrifuged at 10,000 rpm at 4°C and the cell free supernatants were used for the extractions of secondary metabolites. Each culture supernatant was extracted three times by manual shaking with equal volume of ethyl acetate (1:1) in a separating funnel. The solvent layer was collected and concentrated using a rotary evaporator (Buchi, Switzerland) to obtain crude metabolites. The antiviral activity of ethyl acetate actinomycetes extract against WSSV was determined using standard protocol.

### 2.5. Preliminary Screening of antiviral activity

The antiviral activity of actinomycetes extract against WSSV was tested in shrimp based on the procedures described by Balasubramanian et al. (22, 23). Extracts prepared from different isolates of actinomyctes using ethyl acetate were tested to determine the antiviral activity against WSSV. The inactivation of WSSV was confirmed by bioassay, PCR and Western blot analyses. For screening antiviral activity, the shrimp (5 animals per tank in triplicate) were injected intramuscularly with a mixture of viral suspension, extract of actinomycetes and NTE buffer (0.2 M NaCl, 0.02 M Tris-HCl and 0.02 M EDTA, pH 7.4) at the volume of 30 μl per animal (5 μl of viral suspension, 10 μl of actinomyctes extract with varying concentration {100 or 500 μg/animal} and 15 μl of NTE buffer). The positive control consisted of a mixture of 25 μl NTE buffer and 5 μl viral suspension. For toxicity studies, the animals were injected with the extracts of actinomycetes of various concentrations as mentioned above. All these mixtures were incubated at 29 °C for 30 min to 1 h before injection. After incubation, the mixture was injected into experimental animals intramuscularly. Mortalities were recorded for each day of experimental period after post infection with WSSV.

### 2.6. Characteristics of potential isolate

Morphological and cultural characteristics and pigment production of actinomycetes isolate having antiviral activity were examined as described by Shirling and Gottlieb (24). Morphological characters such as colony characteristics, pigment production, absence or presence of aerial and substrate mycelium were observed on actinomycetes isolate grown on International Streptomyces Project (ISP) media 1 to 8. The arrangement of spores and sporulating structures were examined microscopically by using cover slip culture method (25, 26). The spore morphology was studied by scanning electron microscope using electron grid at different magnifications. Cell wall amino acids and whole cell sugars of actinomycetes isolate were analyzed as per the standard protocols (27, 28). The growth of the potential isolate was studied under various conditions such as temperature which ranged from 15 to 40°C, pH from 4 to 10 and NaCl concentrations from 0 to 10% to optimize the growth conditions using standard protocols. For starch utilization test, actinomycetes isolate was streaked on starch agar plate and incubated at 28°C for 7 d. After incubation, iodine solution was poured on the agar and examined for the hydrolysis of starch by the production of clear zone around the bacterial growth. Potential actinomycetes isolate was streaked on gelatin agar plates and incubated at 28°C for 7 d for testing gelatin hydrolysis. After incubation, plates were flooded with 1 mL of mercuric chloride solution and observed for zone of hydrolysis.

To test the casein hydrolysis, isolate was streaked on skimmed milk agar medium and incubated at 28°C for 7 d and observed for zone of hydrolysis. In the urea hydrolysis test, isolates were inoculated into sterile urea agar slants and incubated at 28°C for 7 d and change in colour was observed. Isolates were inoculated into Simon’s citrate slant agar and incubated at 28°C for 7 d and a change in colour was observed. In addition to above tests, other biochemical characters were studied using Ready prepared KB003 Hi 25 kit (HiMedia, India). The tests were ONPG, Lysine, Ornithine, Urease, Phenylalanine, Nitrate reduction, H_2_S production, Citrate utilization, Voges Proskauer’s, Methyl red test, Indole production, Malonate, Esculin hydrolysis, Oxidase, Catalase, Crystal violet, Tween – 20, Sodium azade (0.01%), Sodium azade (0.02%), growth in Macconkey agar and Casein utilization. All reaction wells of the kit (Strip-I and Strip-II) were inoculated with a loop of actinomycetes culture and incubated at 28°C for 14 days and the results were observed.

The ability of actinomycetes to utilize various amino acids as energy source was tested by using basal medium with 0.1% of various amino acids such as L-Arginine, L-Phenyl alanine, L-Tyrosine, L-Methionine, L-Histidine, DL-Tryptophane, L-Cysteine and L-Glutamine. The ability of actinomycetes isolates to utilize various carbon compounds as energy source was tested using ready prepared HiMedia kit (KB009 HiCarbo^™^ kit KB009A /KB009B1 /KB009C). The kit has 35 sugars like Lactose, Xylose, Maltose, Fructose, Dextrose, Galactose, Raffinose, Trehalose, Melibiose, Sucrose, L-Arabinose, Mannose, Inulin, Sodium gluconate, Glycerol, Salicin, Dulcitol, Inositol, Sorbitol, Mannitol, Adonitol, Arabitol, Erythritol, alpha-Methyl-D-glucoside, Rhamnose, Cellobiose, Melezitose, alpha-Methyl-D-Mannoside, Xylitol, ONPG, Esculin, D-Arabinose, Citrate, Malona and Sorbose, and 1 control. The wells in kits (Part A, Part B1 and Part C) were inoculated with actinomycetes isolates and incubated at 28°C for 14 days. After incubation, the wells in the kits were observed for bacterial growth by comparing with negative and positive controls.

The genomic DNA was extracted from actinomycetes isolate using the phenol-chloroform method and 16S rDNA gene was amplified using the universal primer (5′ AGA GTT TGA TCM TGG CTC AG 3′ and 5′ TAC GGY TAC CTT GTT ACG ACT T 3′) with the standard PCR protocol. The PCR product was purified using QIAquick PCR Purification Kit (Qiagen, India) and sequenced using an ABI3730xl sequencer. The nucleotides of the 16S rDNA sequence was matched with the other microbes in the NCBI database using the BLAST program. The sequence was deposited in GenBank. The construction of phylogenetic tree was carried out by using Geneious Basic software and evolutionary history inferred using the neighbor joining method (29).

### 2.7. Purification of bioactive compound

The purification of the bioactive compound having strong antiviral activity against WSSV was carried out using silica gel column chromatography as described by Atta et al. (2009). Twenty gram of silica gel (60 – 120 mm mesh) was mixed with 5 g of ethyl acetate extract of actinomycetes using absolute chloroform and kept for a few hours for drying. Then the mixture was packed in 50 × 2.5 cm size column. The sample was eluted with the petroleum ether and ethyl acetate solvents in the ratio of 60:40 at the flow rate of one drop per minute. All eluted fractions were concentrated at room temperature and checked for antiviral activity against WSSV using the protocol described above. The active fraction (F1) was further fractioned using column chromatography using petroleum ether and methanol in the ratio of 70:30 as a mobile phase. Sub-fractions (F1A and F1B) were collected as described above and determined the antiviral activity.

### 2.8. Identification of the active compound

The compounds obtained from potential actinomycetes isolates were dissolved in ethyl acetate at a concentration of 100 mg/ml and the ultraviolet spectra were recorded on Shimadzu UV - Vis 2450 Spectrophotometer at 200 - 400 nm range using UV-Probe software. A 13 mm Potassium bromide (KBr) pellet using active compounds was prepared and subjected to FTIR analysis to obtain the spectrum between 400–4000 cm^−1^ at a resolution of 4 cm^−1^ using Shimadzu IR affinity–1 FTIR spectrometer (Japan). The purified bioactive compounds isolated from potential actinomycetes isolate were analyzed by using GC Clarus 500 (Perkin Elmer, Singapore) equipped with an Elite–5MS column (30.0 m × 250 μm× 0.25 μm composed of 5% diphenyl–and 95% dimethyl–polysiloxane). The detection of the compound was based on 90% similarity between the MS spectra of the unknown compound and reference compounds accessible in the MS spectra library of National Institute for Standards and Technology (NIST). The ^1^H and ^13^C NMR spectra for the purified compounds having antiviral activity against WSSV were recorded on spectrometer (Bruker, Germany, 400 MHz) using CDCl3 as the solvent and tetramethylsilane as the internal reference. The structure of the antiviral compound extracted from *Streptomyces ghanaensis* CAHSH2 was established with the help of spectral data obtained from spectroscopic analysis. The 2D structure of the compound was obtained using Chem3D Draw Ultra software (Version 10) (30).

### 2.9. Molecular docking studies

3-Dimensional (3D) structures of structural proteins VP26 (PDB id:2edm) and VP28 (PDB id: 2ed6) of WSSV were obtained from Protein Data Bank (PDB) (Berman, *et al*., 2000). The 3D coordinates of VP26 and VP28 were imported and visualized in PyMOL viewer (31). From PDB coordinates, co-crystallized ligand atoms were identified and removed. Then crystallographic water molecules eliminated from the viral protein coordinates and hydrogen atoms were added to the 3D structure and energy minimizations were performed using Swiss pdb viewer (32). The structure of antiviral compounds (di-n-octyl phthalate and bis (2-methylheptyl] phthalate) were drawn in Chemsketch (33) and 3D coordinates were generated using Corina online 3D conversion software (34). These two compounds were then optimized using chimera for flexible conformations (35). The molecular structure-based docking studies were carried out using Auto-Dock version 4.0 (36). These two antiviral compounds were then docked to the two viral proteins, VP26 and VP28. Lamarckian genetic docking algorithm was utilized to generate accurate binding conformations of di-n-octyl phthalate and bis (2-methylheptyl) phthalate in VP26 and VP28 (37).

### 2.10. Confirmation of antiviral activity of EtOAcE against WSSV

The antiviral activity of ethyl acetate extract of actinomyctes isolate (EtOAcE) against WSSV of shrimp was confirmed by polymerase chain reaction (PCR), reverse transcriptase – PCR (RT-PCR), Western blot, ELISA and survival of shrimp in bioassay using standard protocols (38, 39, 40, 41). Quantitative real-time PCR was performed to determine the WSSV viral load in gill tissue of shrimp injected with EtOAcE-treated WSSV and other shrimp groups. The viral copy number was estimated by Step One Plus Real-Time PCR System (Applied Biosystems, USA) using Taqman assay (42).

### 2.11. Toxicity studies

The toxicity of ethyl acetate extract of potential isolate and pure compound having strong antiviral activity against WSSV was evaluated in *Artemia* nauplii and post-larvae of *L. vannamei* by bathing treatment and in adult by intramuscular injection using standard protocols.

## 3. Results

### 3.1. Preliminary screening of actinomycetes isolates against WSSV

Sixty-four actinomycetes isolates collected from different environments were screened for antiviral activity against WSSV. Ethyl acetate extracts of five isolates showed antiviral activity against WSSV at the concentration of 1 mg per shrimp. One isolate designated as CAHSH-2 showed strong antiviral activity and inactivated WSSV at the concentration of 0.2 mg per shrimp and more than 90% survival was observed whereas other isolates failed to inactivate WSSV at lower concentrations. The CAHSH-2 isolate was isolated from plant and selected for detailed studies. The laboratory trial of determining antiviral activity of ethyl extract of CAHSH-2 against WSSV was carried out 21 times since 2014, and the results are given in Table 1. No survival of shrimp injected with WSSV was observed at 48 h p.i. The survival percentage ranged from 88.8 to 98 in shrimp injected with NTE buffer. In the case of shrimp injected with CAHSH-2-extarct treated WSSV, the survival was found to be high and ranged from 88.8 to 100%. The shrimp injected with the extract of CAHSH-2 only showed survival which ranged from 81.4 to 98.3%.

**Table 1.** Date wise laboratory trials carried out to assess the antiviral activity of EtOAcE of Actinomycetes isolate (CAHSH-2) against WSSV since 2014.

| Date of experiment<sup>&</sup> | No of animals used | Average survival of negative control<sup>*</sup> | Average survival of positive control<sup>@</sup> | Average survival of shrimp injected with WSSV treated with EtOAcE<sup>\$</sup> | Average survival of shrimp injected with EtOAcE only<sup>#</sup> |
| --- | --- | --- | --- | --- | --- |
| 12.07.2014 | 60 | 98.3±2.14 | 0 | 96.6±3.16 | 98.3±3.42 |
| 13.08.2014 | 24 | 96.0±2.06 | 0 | 96.0±3.05 | 92.0±2.08 |
| 15.08.2014 | 33 | 97.1±3.15 | 0 | 94.2±2.26 | 97.1±1.96 |
| 19.08.2014 | 21 | 96.6±2.48 | 0 | 95.0±2.57 | 90.0±2.88 |
| 10.09.2014 | 30 | 96.6±3.33 | 0 | 93.3±3.33 | 96.6±3.66 |
| 01.10.2014 | 42 | 97.5±2.29 | 0 | 95.0±2.19 | 92.5±2.49 |
| 26.02.2015 | 24 | 96.0±2.61 | 0 | 96.0±2.99 | 92.0±4.15 |
| 01.03.2015 | 27 | 96.2±3.32 | 0 | 92.5±1.87 | 96.2±3.84 |
| 04.04.2015 | 15 | 93.3±3.38 | 0 | 100 | 86.6±6.66 |
| 08.04.2015 | 18 | 94.4±2.28 | 0 | 88.8±2.15 | 94.4±2.16 |
| 25.07.2015 | 48 | 98.0±2.50 | 0 | 96.0±3.26 | 94.0±2.95 |
| 06.08.2015 | 30 | 96.6±3.33 | 0 | 93.3±5.12 | 96.6±3.33 |
| 04.09.2015 | 24 | 96.0±4.61 | 0 | 96.0±3.30 | 92.0±2.49 |
| 21.09.2015 | 21 | 96.6±3.33 | 0 | 95.0±2.14 | 90.0±2.09 |
| 21.11.2015 | 33 | 97.1±2.19 | 0 | 94.2±2.55 | 97.1±3.19 |
| 08.01.2016 | 48 | 98.0±2.49 | 0 | 96.0±3.66 | 94.0±4.08 |
| 04.02.2016 | 15 | 93.3±3.33 | 0 | 93.3±3.33 | 86.6±3.33 |
| 10.02.2016 | 27 | 96.2±2.16 | 0 | 100 | 96.2±2.49 |
| 03.03.2016 | 18 | 88.8±2.28 | 0 | 94.4±2.27 | 88.8±2.26 |
| 14.03.2016 | 30 | 93.3±3.33 | 0 | 96.3±2.28 | 93.3±3.34 |
| 25.05.2016 | 27 | 96.2±2.13 | 0 | 92.5±3.19 | 81.4±2.83 |

\*Shrimp injected with NTE buffer; <sup>@</sup>Shrimp injected with WSSV (5 µl of haemolymph from moribund WSSV-infected shrimp per shrimp) intramuscularly; <sup>\$</sup>Shrimp injected with mixture of WSSV (5 µl of WSSV inoculum) and Actinomycetes extract at the concentration of 200 µg per shrimp; <sup>#</sup>Shrimp injected with actinomycetes extract only. <sup>&</sup>Experimental period was 20 days.

### 3.2. Characteristics of potential Actinomycetes isolates

The CAHSH-2 isolate having strong antiviral activity against WSSV was characterized by colony morphology, spore structure, cultural characteristics, physiological tests, biochemical tests, carbon utilization, nitrogen utilization and antibiotic sensitivity, and molecular taxonomy by sequencing 16S rRNA gene. The results are presented in table 2 and Fig.1. All these characteristics of isolates were studied along with reference strains, *Streptomyces ghanaensis* and *S. viridosporus* obtained from IMTECH, Chandigarh, India. The colony of CAHSH-2 contained both aerial and substrate mycelia. Microscopic observation of actinobacterial isolate revealed the presence of well-developed short chain and branched mycelium which consisted of verticillate type spores (Fig 2). The isolated CAHSH-2 strain is a Gram positive and non-motile actinomycete. The optimal growth conditions of the isolate were temperature at 28 °C, pH of 7 and NaCl concentration of 2%. This isolate utilized a wide variety of carbohydrates as a carbon source. Histidine among various amino acids tested was found to be the best source of nitrogen for the growth of CAHSH-2. The potential isolate utilized lysine and ornithine but failed to deaminate phenylalanine. This isolate failed to reduce the nitrate and did not produce H2S and was found to be positive for urease and citrate utilization showing their ability to hydrolyze urea and citrate. It was found to be positive for VP test, ONPG test, oxidase and MR test, and negative for indole production and catalase. Analysis of whole-cell components revealed LL-type of diaminopimelic acid and glycine, and no characteristic sugar pattern. PCR amplification of 16S rDNA of CAHSH-2 yielded 1311 nucleotides and its sequence was deposited in GenBank under the accession number KT006906.1. The BLAST analysis of the sequence revealed that the CAHSH-2 showed 99% similarity with *Streptomyces ghanaensis* (NR-112460.1). Furthermore, phylogenetic tree was constructed with CAHSH-2 isolate and neighbor joining phylogenetic tree using cluster W software which revealed that it was *Streptomyces* sp. with high bootstrap value (Fig 3). Based on this, CAHSH-2 isolated from *Ocimum tenuiflorum* was identified as *Streptomyces ghanaensis* like strain and designated as *Streptomyces ghanaensis* – CAHSH-2

**Fig.1.**
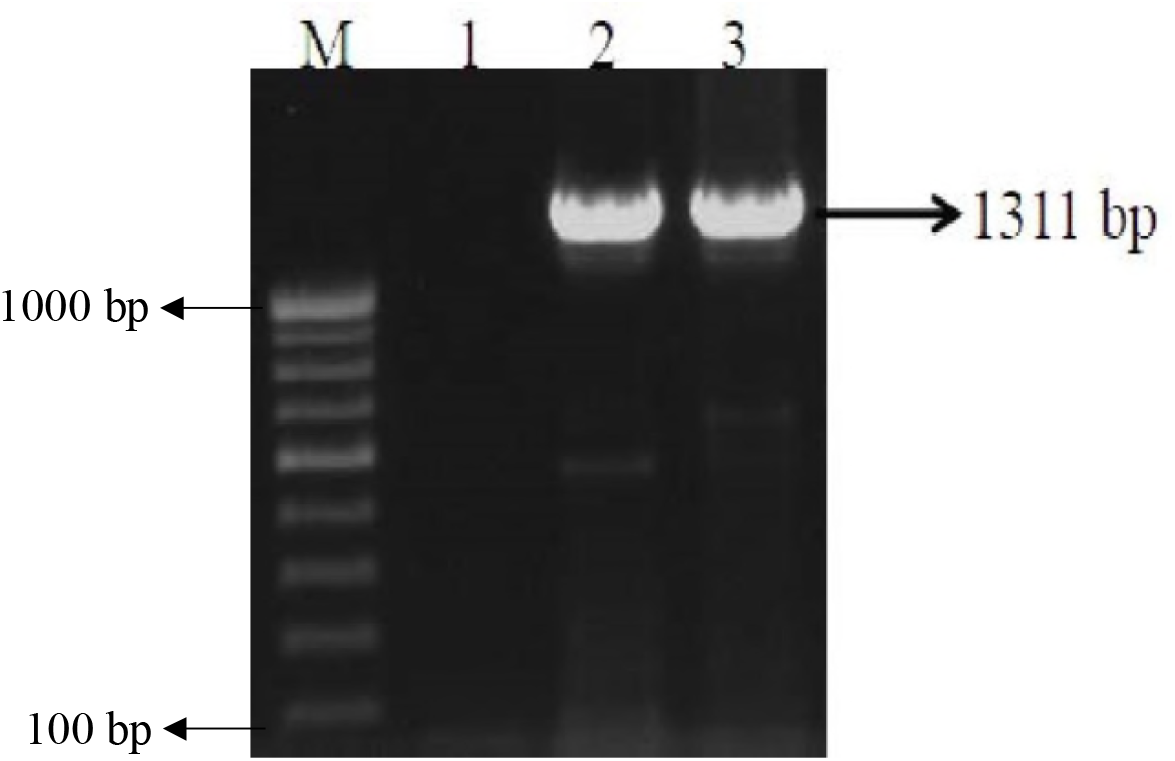
Amplification of 16S rRNA gene of Actinomycetes isolate CAHSH-2 and reference strain of *Streptomyces ghanaensis* obtained from IMTECH. Lane M-100 bp Marker; Lane 1-Negative Control; Lane 2-CAHSH-2; Lane 3-*Streptomyces ghanaensis*

**Fig. 2.**
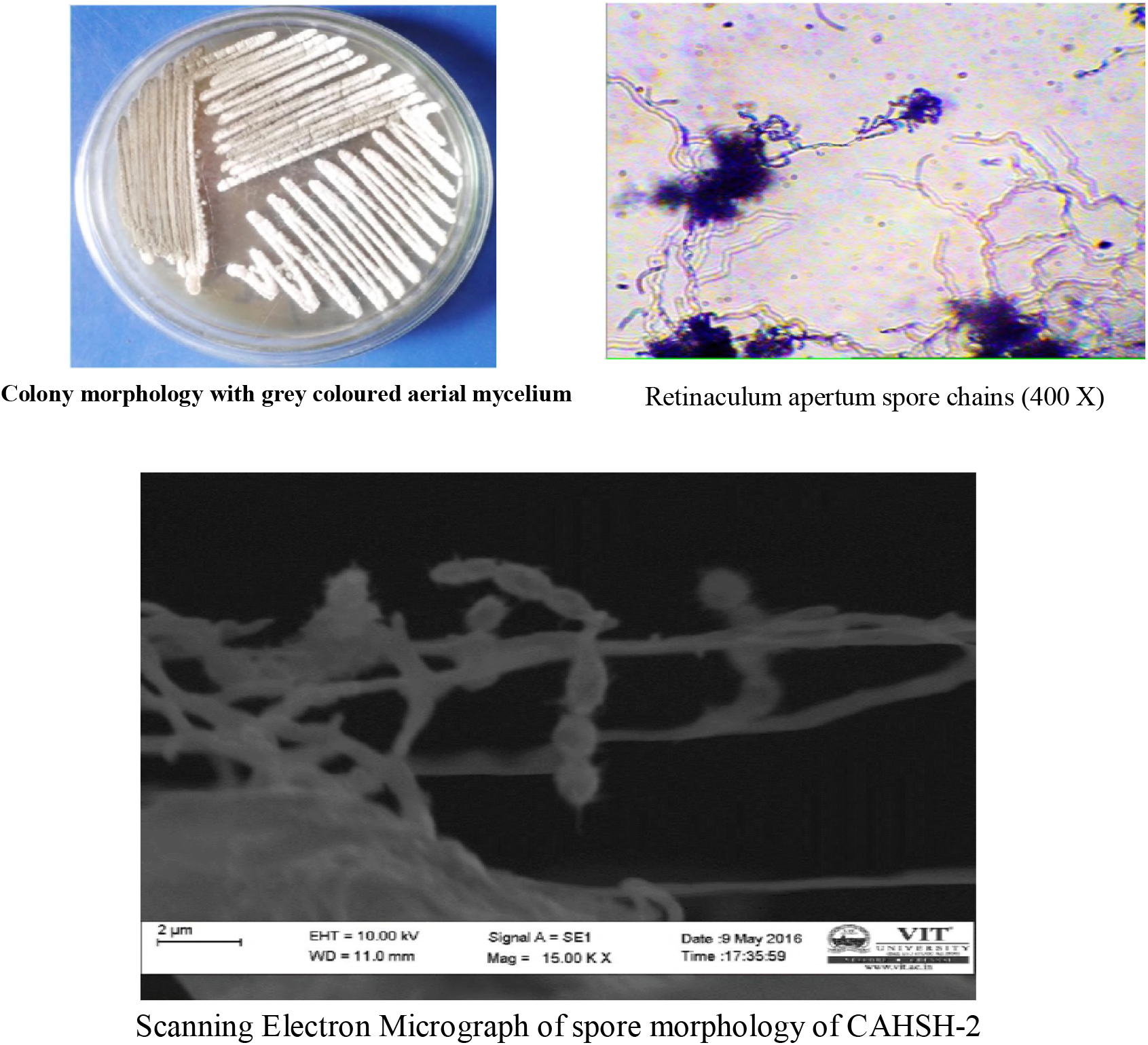
Morphological characteristics of CAHSH-2 after 14 days of growth at 28 ± 2°C on ISP medium-3

**Fig. 3.**
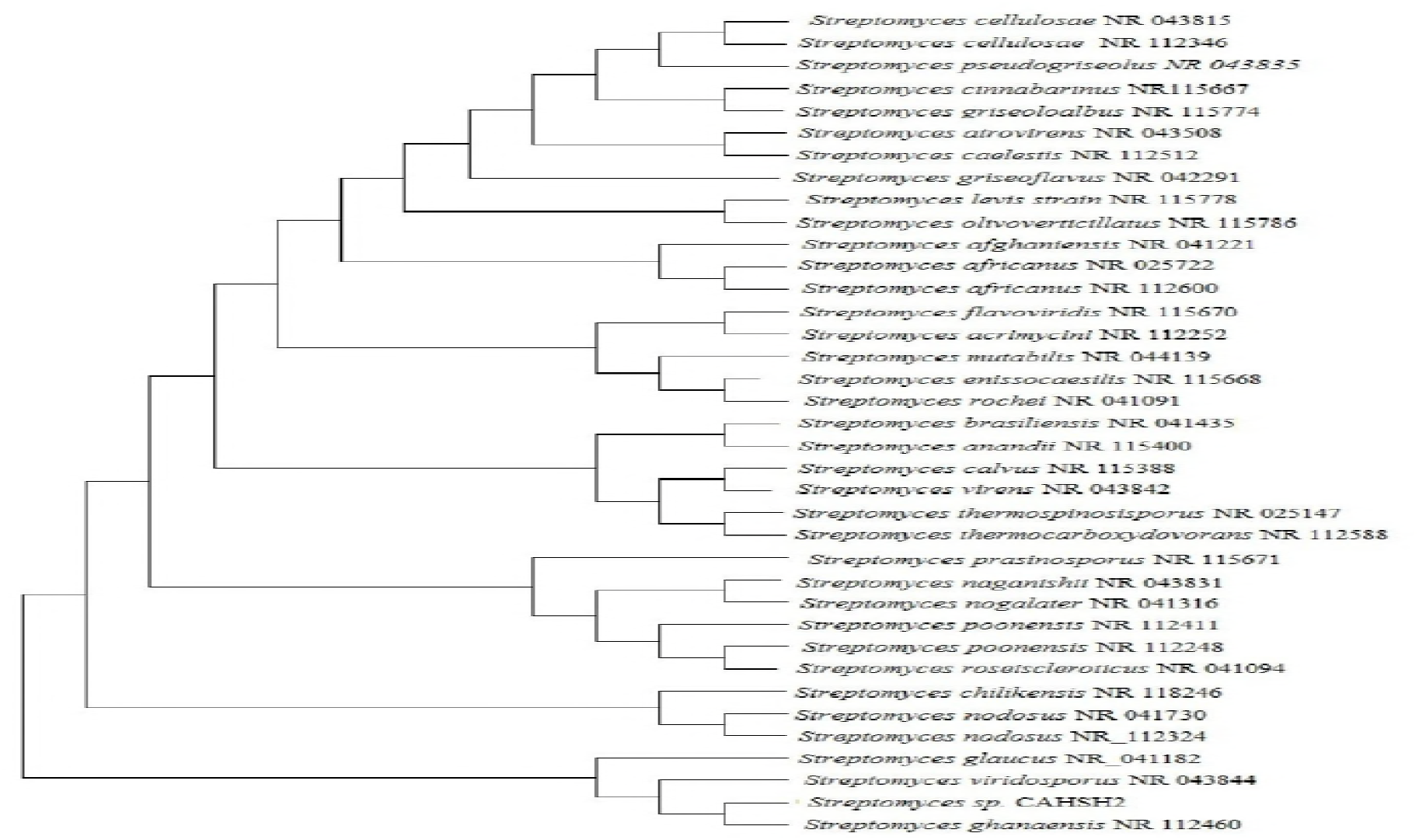
Rooted neighbour joining phylogenetic tree of actinomycetes CAHSH-2 based on 16S rRNA gene sequences, showing the relationship between actinomycetes CAHSH-2 and related representative species of the *Streptomye ghanaensis*.

**Table 2a.** Morphological characteristics of CAHSH-2 and CAHSH-3 along with reference strains from IMTECH

| Characteristics | Actinomycetes CAHSH-2 |  |  | Actinomycetes CAHSH -3 |  |  | Reference strain |  |
| --- | --- | --- | --- | --- | --- | --- | --- | --- |
|  | IS1 | IS2 | IS3 | IS1 | IS2 | IS3 | *Streptomyces ghanaensis* | *Streptomyces viridosporus* |
| Spore morphological characteristics | +++ | +++ | +++ | +++ | +++ | +++ | +++ | +++ |
| Presence of Aerial mycelium | +++ | +++ | +++ | +++ | +++ | +++ | +++ | +++ |
| Presence of Substrate mycelium | Dark grey | Dark grey | Dark grey | Green | Green | Green | Dark grey | Green |
| Colour of aerial mycelium | Yellow | Yellow | Yellow | Cream | Cream | Cream | Yellow | Cream |
| Colour of substrate mycelium | Branched | Branched | Branched | Branched & twisted | Branched & twisted | Branched & twisted | Branched | Branched & twisted |
| Spore type | Retinaculum apertum | Retinaculum apertum | Retinaculum apertum | Spiral | Spiral | Spiral | Retinaculum apertum | Spiral |
| Spore shape | Oval | Oval | Oval | Oval | Oval | Oval | Oval | Oval |
| Spore Surface | Thorny | Thorny | Thorny | Spiny | Spiny | Spiny | Thorny | Spiny |
| Spore chain | Long chain | Long chain | Long chain | Long chain | Long chain | Long chain | Long chain | Long chain |
| Spore motility | Non-motile | Non-motile | Non-motile | Non-motile | Non-motile | Non-motile | Non-motile | Non-motile |
| Acid fast | + | + | + | + | + | + | + | + |
| Sporangia | - | - | - | - | - | - | - | - |
| Melanin production | Green | Green | Green | - | - | - | Green | - |
| Diffusible pigments | + ve | + ve | + ve | + ve | + ve | + ve | + ve | + ve |

IS – in triplicate

**Table 2b.**
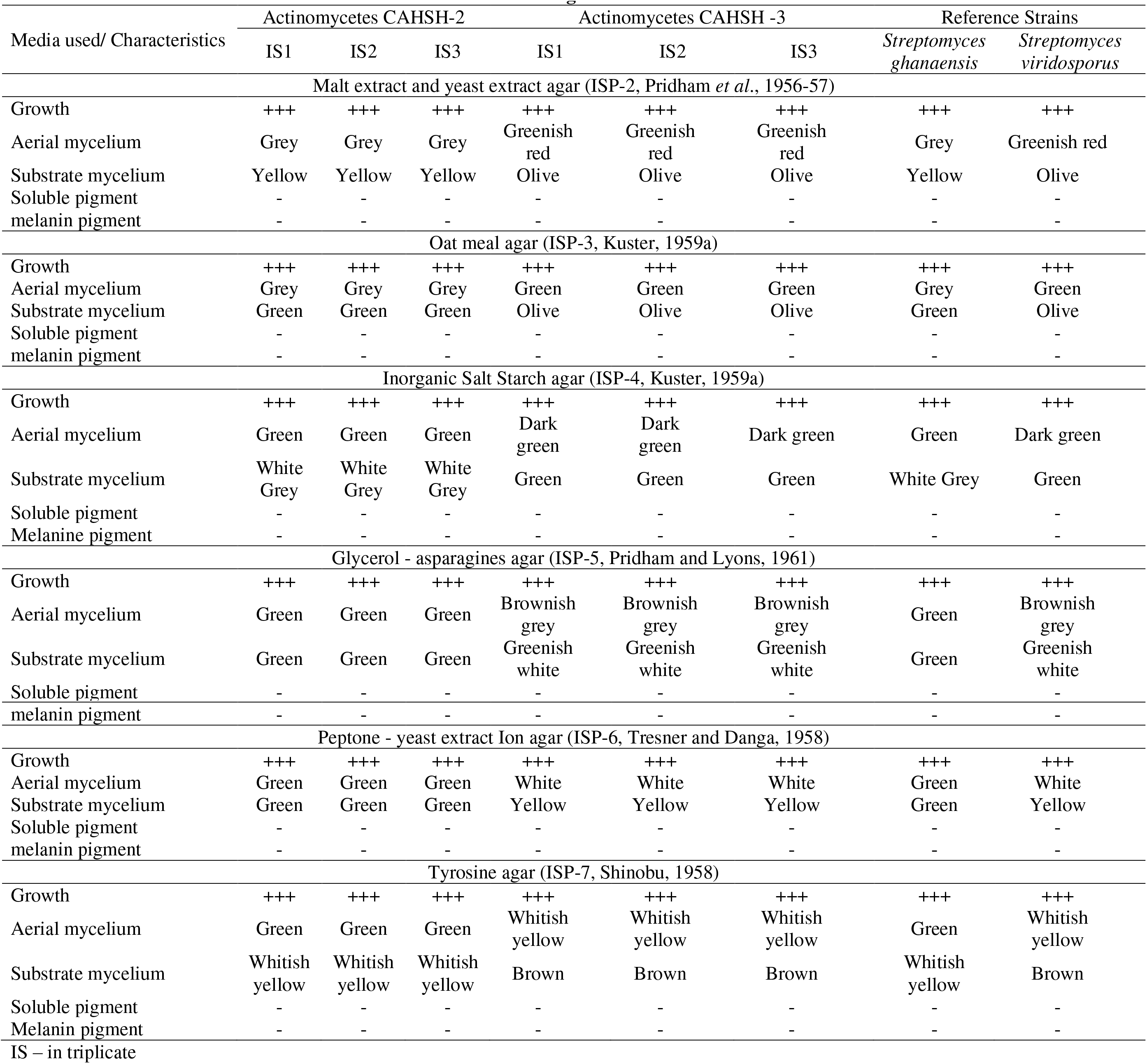
Cultural characteristics of CAHSH-2 and CAHSH-3 along with reference strains from IMTECH

**Table 2c.**
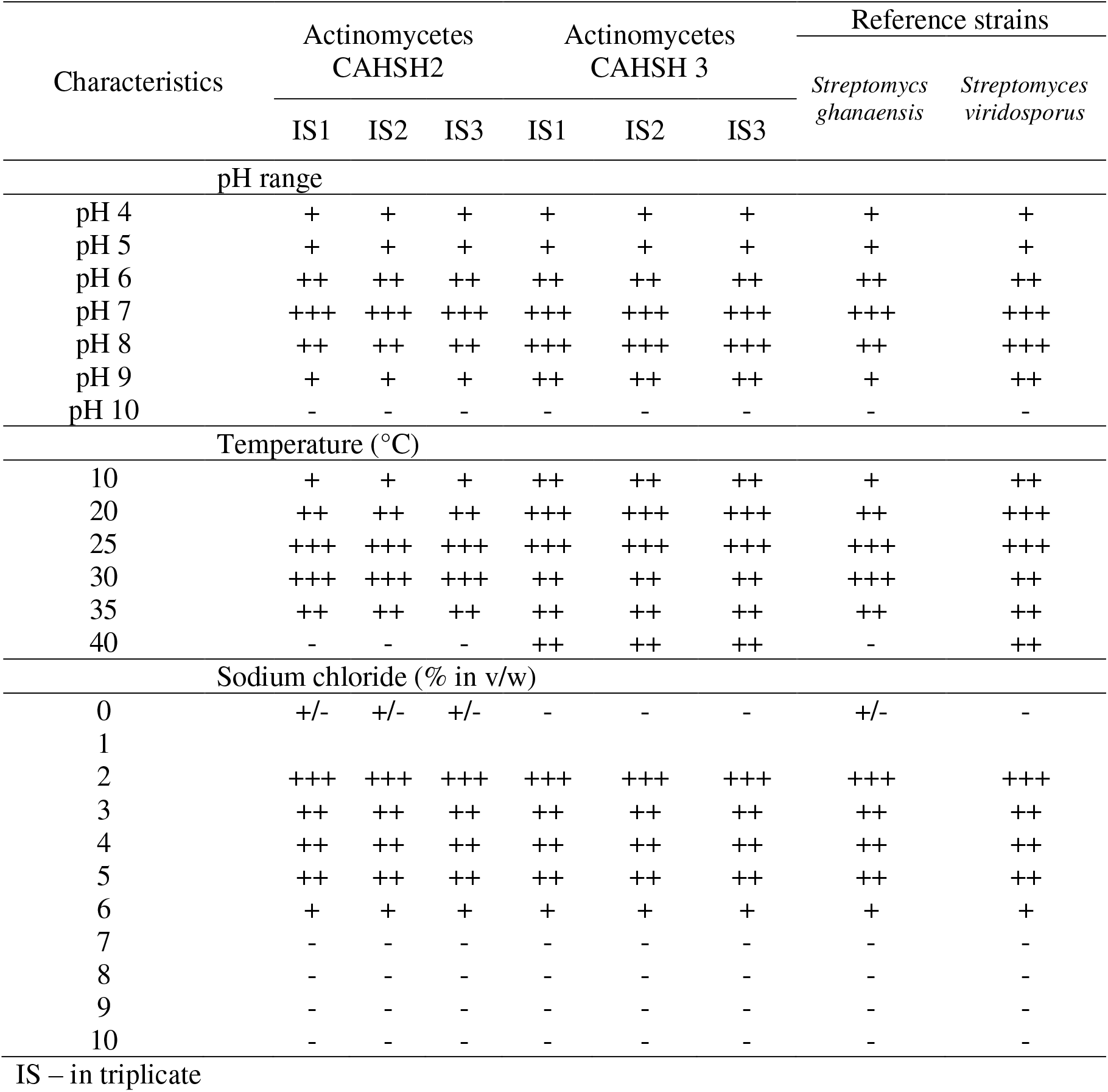
Physiological characteristics of CAHSH-2 and CAHSH-3 along with reference strains from IMTECH

| Characteristics | Actinomycetes<br>CAHSH2 |  |  | Actinomycetes<br>CAHSH 3 |  |  | Reference strains |  |
| --- | --- | --- | --- | --- | --- | --- | --- | --- |
|  | IS1 | IS2 | IS3 | IS1 | IS2 | IS3 | *Streptomyces*<br>*ghanaensis* | *Streptomyces*<br>*viridosporus* |
| pH range |  |  |  |  |  |  |  |  |
| pH 4 | + | + | + | + | + | + | + | + |
| pH 5 | + | + | + | + | + | + | + | + |
| pH 6 | ++ | ++ | ++ | ++ | ++ | ++ | ++ | ++ |
| pH 7 | +++ | +++ | +++ | +++ | +++ | +++ | +++ | +++ |
| pH 8 | ++ | ++ | ++ | +++ | +++ | +++ | ++ | +++ |
| pH 9 | + | + | + | ++ | ++ | ++ | + | ++ |
| pH 10 | - | - | - | - | - | - | - | - |
| Temperature (°C) |  |  |  |  |  |  |  |  |
| 10 | + | + | + | ++ | ++ | ++ | + | ++ |
| 20 | ++ | ++ | ++ | +++ | +++ | +++ | ++ | +++ |
| 25 | +++ | +++ | +++ | +++ | +++ | +++ | +++ | +++ |
| 30 | +++ | +++ | +++ | ++ | ++ | ++ | +++ | ++ |
| 35 | ++ | ++ | ++ | ++ | ++ | ++ | ++ | ++ |
| 40 | - | - | - | ++ | ++ | ++ | - | ++ |
| Sodium chloride (% in v/w) |  |  |  |  |  |  |  |  |
| 0 | +/- | +/- | +/- | - | - | - | +/- | - |
| 1 |  |  |  |  |  |  |  |  |
| 2 | +++ | +++ | +++ | +++ | +++ | +++ | +++ | +++ |
| 3 | ++ | ++ | ++ | ++ | ++ | ++ | ++ | ++ |
| 4 | ++ | ++ | ++ | ++ | ++ | ++ | ++ | ++ |
| 5 | ++ | ++ | ++ | ++ | ++ | ++ | ++ | ++ |
| 6 | + | + | + | + | + | + | + | + |
| 7 | - | - | - | - | - | - | - | - |
| 8 | - | - | - | - | - | - | - | - |
| 9 | - | - | - | - | - | - | - | - |
| 10 | - | - | - | - | - | - | - | - |

IS – in triplicate

**Table 2d.**
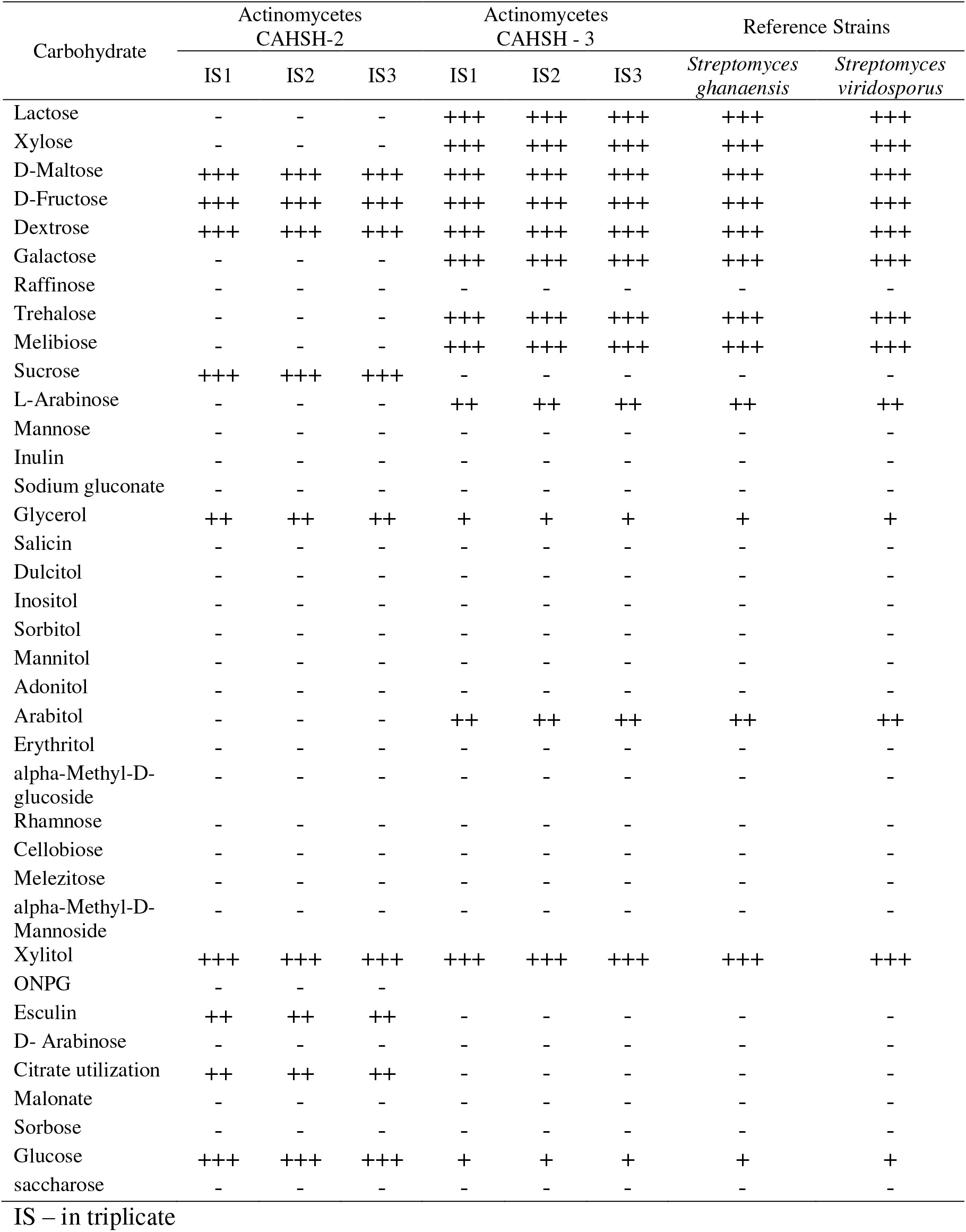
Carbohydrate utilization test forCAHSH-2 and CAHSH-3 along with reference strains from IMTECH.

**Table 2e.**
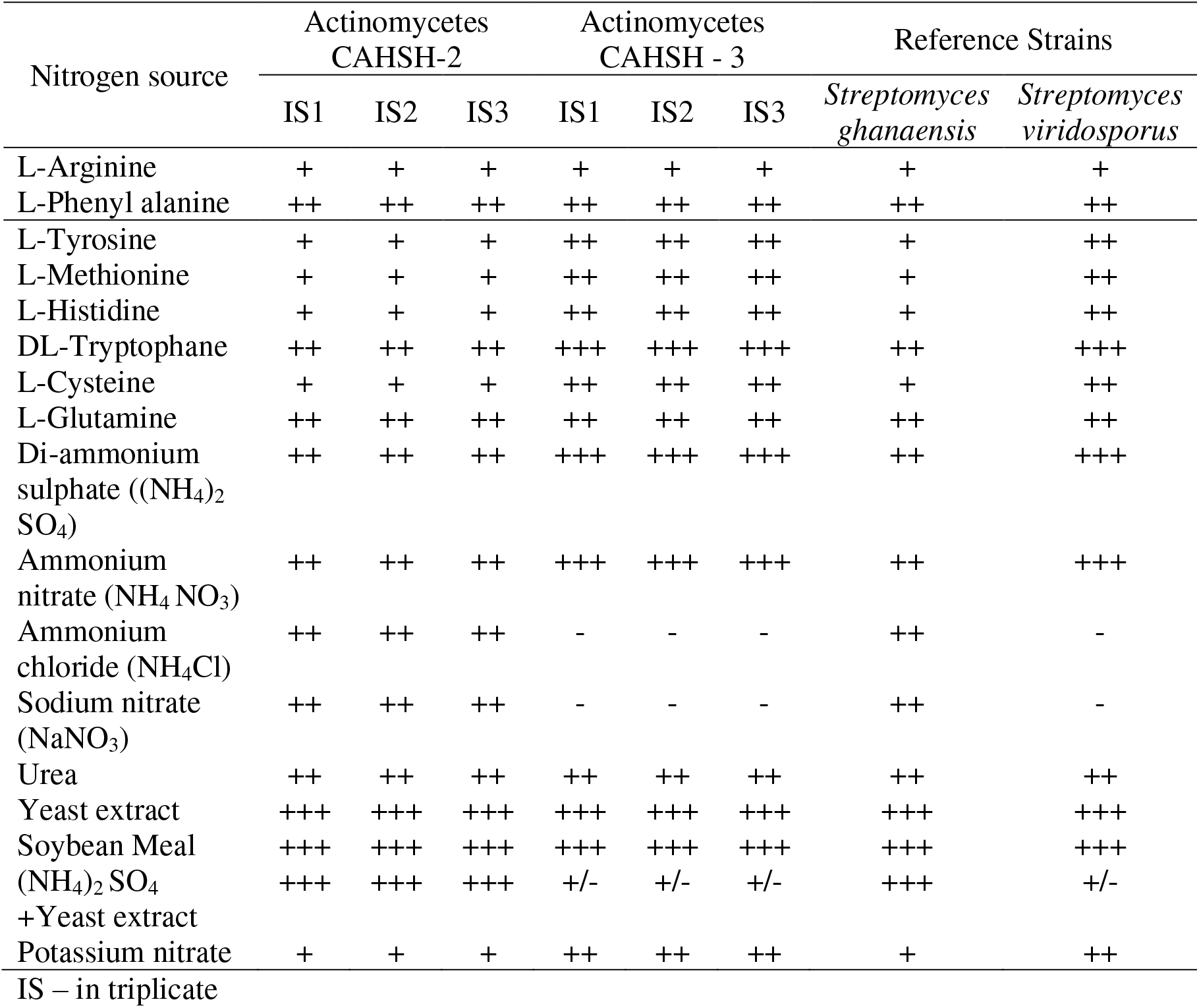
Amino acid and nitrogen utilization tests for CAHSH-2 and CAHSH-3 along with reference strains from IMTECH.

**Table 2f.**
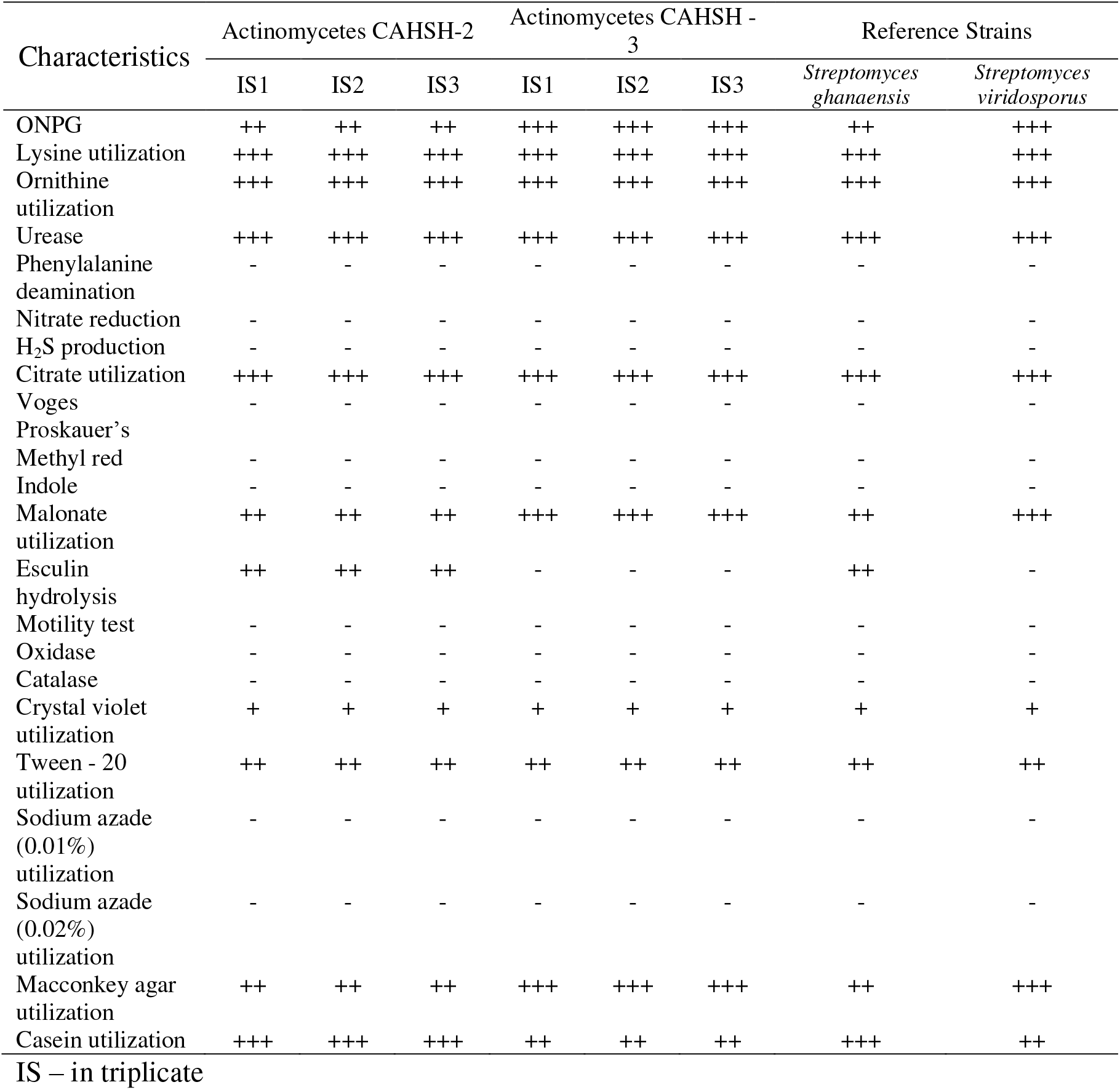
Biochemical characterizations of CAHSH-2 and CAHSH-3 along with reference strains from IMTECH.

**Table 2g.** Chemotaxonomy characterizations of CAHSH-2 and CAHSH-3 along with reference strains from IMTECH (IS – in triplicate; I - Pattern I).

| Characteristics | Actinomycetes CAHSH-2 |  |  | Actinomycetes CAHSH - 3 |  |  | Reference Strains |  |
| --- | --- | --- | --- | --- | --- | --- | --- | --- |
|  | IS1 | IS2 | IS3 | IS1 | IS2 | IS3 | *Streptomyces ghanaensis* | *Streptomyces viridosporus* |
| Cell wall amino acid | LL-DAP with glycine | LL-DAP with glycine | LL-DAP with glycine | LL-DAP with glycine | LL-DAP with glycine | LL-DAP with glycine | LL-DAP with glycine | LL-DAP with glycine |
| Whole cell sugar | No characteristic sugar pattern | No characteristic sugar pattern | No characteristic sugar pattern | No characteristic sugar pattern | No characteristic sugar pattern | No characteristic sugar pattern | No characteristic sugar pattern | No characteristic sugar pattern |
| Cell wall type | I | I | I | I | I | I | I | I |

**Table 2h.** Degradation activities of CAHSH-2 and CAHSH-3 along with reference strains from IMTECH

| Characteristics | Results |  |  |  |  |  | *Streptomyces ghanaensis* | *Streptomyces viridosporus* |
| --- | --- | --- | --- | --- | --- | --- | --- | --- |
|  | CAHSH-2 |  |  | CAHSH-3 |  |  |  |  |
|  | IS1 | IS2 | IS3 | IS1 | IS2 | IS3 |  |  |
| Starch | + | + | + | + | + | + | + | + |
| Casein | + | + | + | + | + | + | + | + |
| L-tyrosine | + | + | + | + | + | + | + | + |
| xylene | - | - | - | + | + | + | - | + |
| xanthine | - | - | - | + | + | + | - | + |

**Table 2i.**
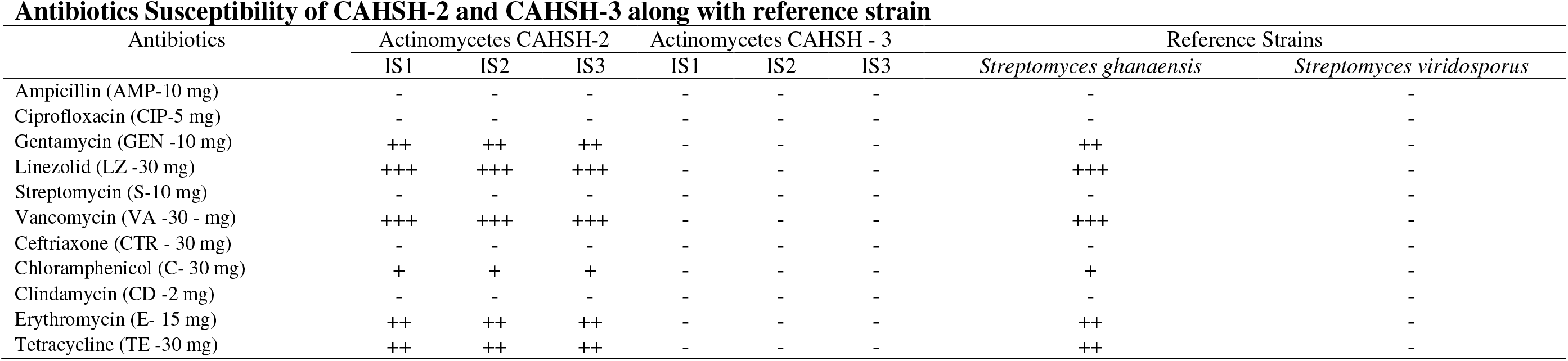
Antibiotics Susceptibility of CAHSH-2 and CAHSH-3 along with reference strain

| Antibiotics | Actinomycetes CAHSH-2 |  |  | Actinomycetes CAHSH - 3 |  |  | Reference Strains |  |
| --- | --- | --- | --- | --- | --- | --- | --- | --- |
|  | IS1 | IS2 | IS3 | IS1 | IS2 | IS3 | *Streptomyces ghanaensis* | *Streptomyces viridosporus* |
| Ampicillin (AMP-10 mg) | - | - | - | - | - | - | - | - |
| Ciprofloxacin (CIP-5 mg) | - | - | - | - | - | - | - | - |
| Gentamycin (GEN -10 mg) | ++ | ++ | ++ | - | - | - | ++ | - |
| Linezolid (LZ -30 mg) | +++ | +++ | +++ | - | - | - | +++ | - |
| Streptomycin (S-10 mg) | - | - | - | - | - | - | - | - |
| Vancomycin (VA -30 - mg) | +++ | +++ | +++ | - | - | - | +++ | - |
| Ceftriaxone (CTR - 30 mg) | - | - | - | - | - | - | - | - |
| Chloramphenicol (C- 30 mg) | + | + | + | - | - | - | + | - |
| Clindamycin (CD -2 mg) | - | - | - | - | - | - | - | - |
| Erythromycin (E- 15 mg) | ++ | ++ | ++ | - | - | - | ++ | - |
| Tetracycline (TE -30 mg) | ++ | ++ | ++ | - | - | - | ++ | - |

### 3.3. Extraction, purification and activity of antiviral compounds

Five prominent bands were observed in TLC plate loaded with ethyl acetate extract having antiviral activity against WSSV (Fig 4). The Rf value of five bands was determined as 0.49-0.62 (Band I); 0.62-0.72 (Band II); 0.72 – 0.87 (Band III) 0.72-0.87 (Band IV) and 0.87-0.92 (Band V). The extract was fractionated by silica gel column and five fractions (F1, F2, F3, F4 and F5) were obtained. Among the five fractions, F1 was found to have strong antiviral activity. The mortality of shrimp injected with F1 treated WSSV at the concentration of 100 μg per shrimp was estimated to be about 0% on 3 d p.i., 5.56% on 5 d p.i and 13.89% on 10 d p.i. The F1 was further fractioned by column chromatography and two sub fractions, F1A and F1B were obtained. These two sub-fractions were tested for their antiviral activity against WSSV and both sub-fractions showed antiviral activity against WSSV (Table 3 a, b). The mortality of shrimp injected with F1B treated WSSV at the concentration of 100 μg per shrimp was estimated to be about 0% on 3 and 5 d p.i., and 11.1% on 10 d p.i. In the case of shrimp injected with F1A treated WSSV, the mortality was found to be 66.7% and 81.5% on 5 and 10 d p.i., respectively at the concentration of 100 μg per shrimp. In the case of shrimp injected with WSSV, 100% mortality was observed on 3 d p.i.

**Fig. 4.**
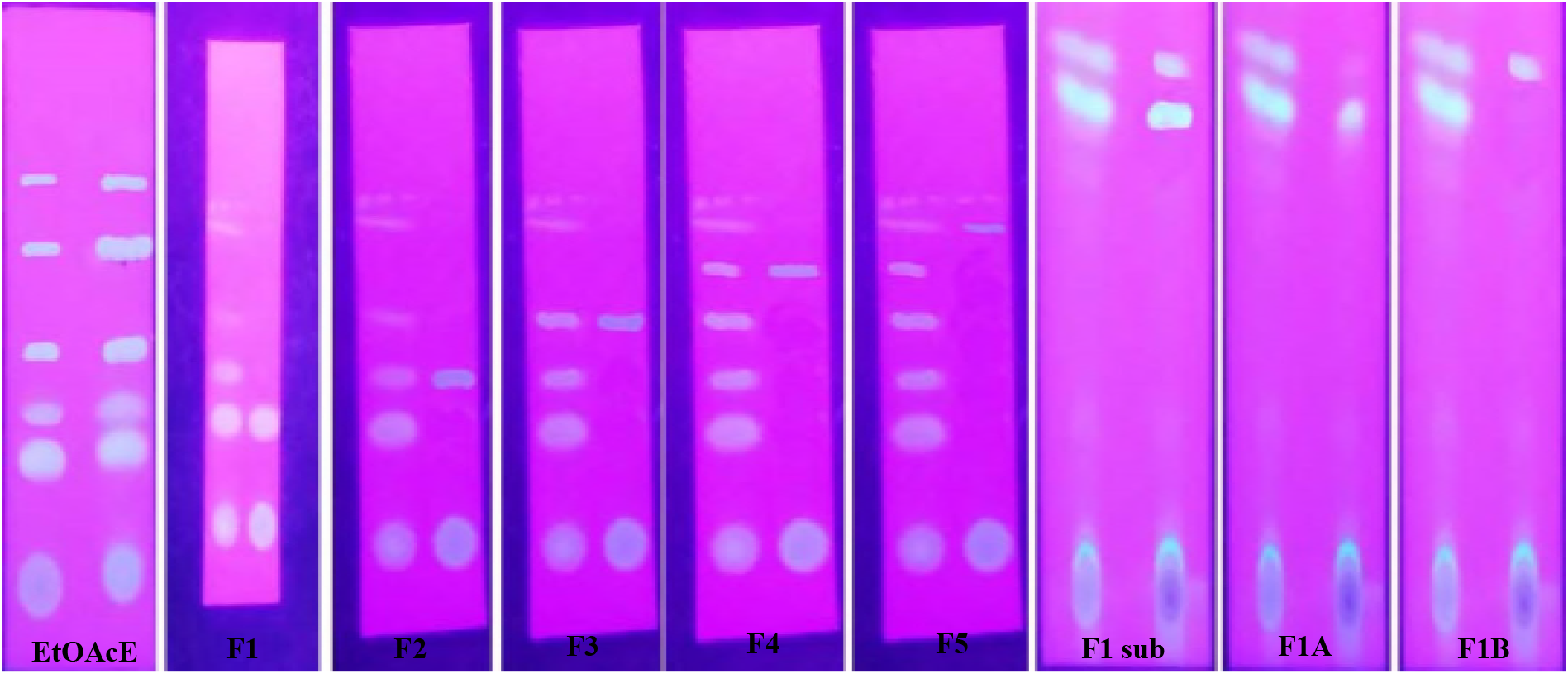
Thin layer chromatographic analysis of EtOAcE of CAHSH-2 and their homogeneity, and TLC analysis of active fraction and its homogeneity.

**Table 3a.** Antiviral activity of different fractions (F1, F2, F3, F4 and F5) from EtOAcE of CAHSH-2 isolate against WSSV in *Litopenaeus vannamei*.

| Experimental Groups | No of shrimp used in each trail<sup>§</sup> 4 trials (n = 117) | Cumulative percentage mortality at different time intervals (days post injection) |  |  |  |
| --- | --- | --- | --- | --- | --- |
|  |  | 2 d | 3 d | 5 d | 10 d |
| Positive control<sup>@</sup> | 3 X 3 = 9 | 83.33±3.33 | 100 | 100 | 100 |
| Negative control* | 3 X 3 = 9 | 0 | 0 | 5.56±0.88 | 11.11±1.15 |
| Extract 200 µg per shrimp<sup>#</sup> | 3 X 3 = 9 | 0 | 0 | 8.33±1.02 | 16.67±1.64 |
| F1 - 100 µg per shrimp** | 3 X 3 = 9 | 0 | 0 | 5.56±0.76 | 13.89±1.25 |
| F1 - 50 µg per shrimp** | 3 X 3 = 9 | 0 | 11.11±0.98 | 19.44±1.16 | 41.67±2.29 |
| F2 - 100 µg per shrimp** | 3 X 3 = 9 | 5.56±0.66 | 16.67±1.15 | 36.11±2.85 | 77.78±3.42 |
| F2 - 50 µg per shrimp** | 3 X 3 = 9 | 11.11±1.05 | 27.78±1.09 | 72.22±2.96 | 94.44±2.77 |
| F3 - 100 µg per shrimp** | 3 X 3 = 9 | 13.89±1.12 | 41.67±2.28 | 80.56±3.51 | 91.67±3.49 |
| F3 - 50 µg per shrimp** | 3 X 3 = 9 | 27.78±1.82 | 58.33±1.94 | 86.11±4.19 | 100 |
| F4 - 100 µg per shrimp** | 3 X 3 = 9 | 36.11±2.82 | 47.22±1.63 | 91.67±3.72 | 100 |
| F4 - 50 µg per shrimp** | 3 X 3 = 9 | 52.78±2.19 | 88.89±2.54 | 100 | 100 |
| F5 - 100 µg per shrimp** | 3 X 3 = 9 | 44.44±2.27 | 86.11±1.14 | 100 | 100 |
| F5 - 50 µg per shrimp** | 3 X 3 = 9 | 55.56±2.73 | 88.89±3.29 | 100 | 100 |

\*Shrimp injected with NTE buffer; <sup>@</sup>Shrimp injected with WSSV (5 µl of haemolymph from moribund WSSV-infected shrimp per shrimp) intramuscularly; \*\*Shrimp injected with mixture of WSSV (5 µl of WSSV inoculum) and different fractions (F1, F2, F3, F4 and F5) from ethyl acetate extract extracted from Actinomyces isolate (CAHSH-2) at different concentrations; <sup>#</sup>Shrimp injected with mixture of WSSV (5 µl of WSSV inoculum) and ethyl acetate extract extracted from Actinomyces isolate (CAHSH-2) at the concentration of 200 µg per shrimp. <sup>§</sup>This trial was carried out for 4 times (21.11.2015, 08.01.2016, 30.01.2016 and 10.02.2016).

**Table 3b.** Antiviral activity of different sub-fractions (F1A and F1B) from separated from Fraction1 of EtOAcE of CAHSH-2 isolate against WSSV in *Litopenaeus vannamei*.

| Experimental Groups | No of shrimp used in each trail<sup>§</sup><br>3 trials<br>(n = 81) | Cumulative percentage mortality at different time intervals<br>(days post injection) |  |  |  |
| --- | --- | --- | --- | --- | --- |
|  |  | 2 d | 3 d | 5 d | 10 d |
| Positive control<sup>@</sup> | 3 X 3 = 9 | 74.07±2.47 | 100 | 100 | 100 |
| Negative control* | 3 X 3 = 9 | 0 | 0 | 7.41±0.68 | 11.11±0.99 |
| Extract 200 µg per shrimp<sup>#</sup> | 3 X 3 = 9 | 0 | 0 | 3.70±0.94 | 11.11±0.85 |
| F1A - 100 µg per shrimp** | 3 X 3 = 9 | 11.11±1.07 | 51.85±1.55 | 66.67±3.14 | 81.48±2.33 |
| F1A - 50 µg per shrimp** | 3 X 3 = 9 | 37.03±1.92 | 77.78±2.18 | 85.18±3.36 | 100 |
| F1A - 25 µg per shrimp** | 3 X 3 = 9 | 70.37±3.35 | 88.89±3.67 | 100 | 100 |
| F1B - 100 µg per shrimp** | 3 X 3 = 9 | 0 | 0 | 0 | 11.11±0.86 |
| F1B - 50 µg per shrimp** | 3 X 3 = 9 | 0 | 11.11±0.75 | 18.51±1.12 | 22.22±1.27 |
| F1B - 25 µg per shrimp** | 3 X 3 = 9 | 0 | 25.92±1.15 | 37.03±1.62 | 66.67±2.37 |

\*Shrimp injected with NTE buffer; <sup>@</sup>Shrimp injected with WSSV (5 µl of haemolymph from moribund WSSV-infected shrimp per shrimp) intramuscularly; \*\*Shrimp injected with mixture of WSSV (5 µl of WSSV inoculum) and different sub-fractions (F1A and F1B) separated from Fraction 1 of ethyl acetate extract extracted from Actinomycetes isolate (CAHSH-2) at different concentrations; <sup>#</sup>Shrimp injected with mixture of WSSV (5 µl of WSSV inoculum) and ethyl acetate extract extracted from Actinomycetes isolate (CAHSH-2) at the concentration of 200 µg per shrimp.
<sup>§</sup>This trial was carried out for 3 times (14.05.2016, 21.05.2016 and 25.06.2016).

### 3.4. Identification of antiviral compounds

The F1A and F1B sub-fractions from active fraction exhibited a maximum UV absorption at 287 nm and 279 nm, respectively in ethyl acetate (Fig 5a). The FTIR spectrum of the F1B showed characteristic peaks corresponding to standard library spectra (Fig 5b). A broad peak observed at 3076.47 cm^−1^ indicated the presence O–H stretching. Peak observed at 2966.38 cm^−1^ indicated aromatic C–H stretching frequency. Peak observed at 1736 cm^−1^ indicated the presence of C=O stretch (ketone). Peak observed at 1639 cm^−1^ corresponded to C=O stretching (esters). Peak at 1462.68 cm^−1^ represented C=C stretching. The FTIR spectrum of F1A showed the absence of hydroxyl group but showed the presence of ester moiety (1714.23 and 1282.90 cm^−1^) and aromatic system (1639.22, 1461.35, 908.70 and 722.32 cm^−1^) (Fig 5b). The results of Gas chromatography - mass spectrum (GC-MS) analysis of antiviral compounds (F1A and F1B) from actinomycetes isolate CAHSH-2 revealed the molecular formula of C_24_H_38_O_4_ with molecular weight of 390 g/mol for F1A and C_24_H_38_O_4_ and 388.54 g/mol for F1B (Fig 5c).

**Fig. 5a.**
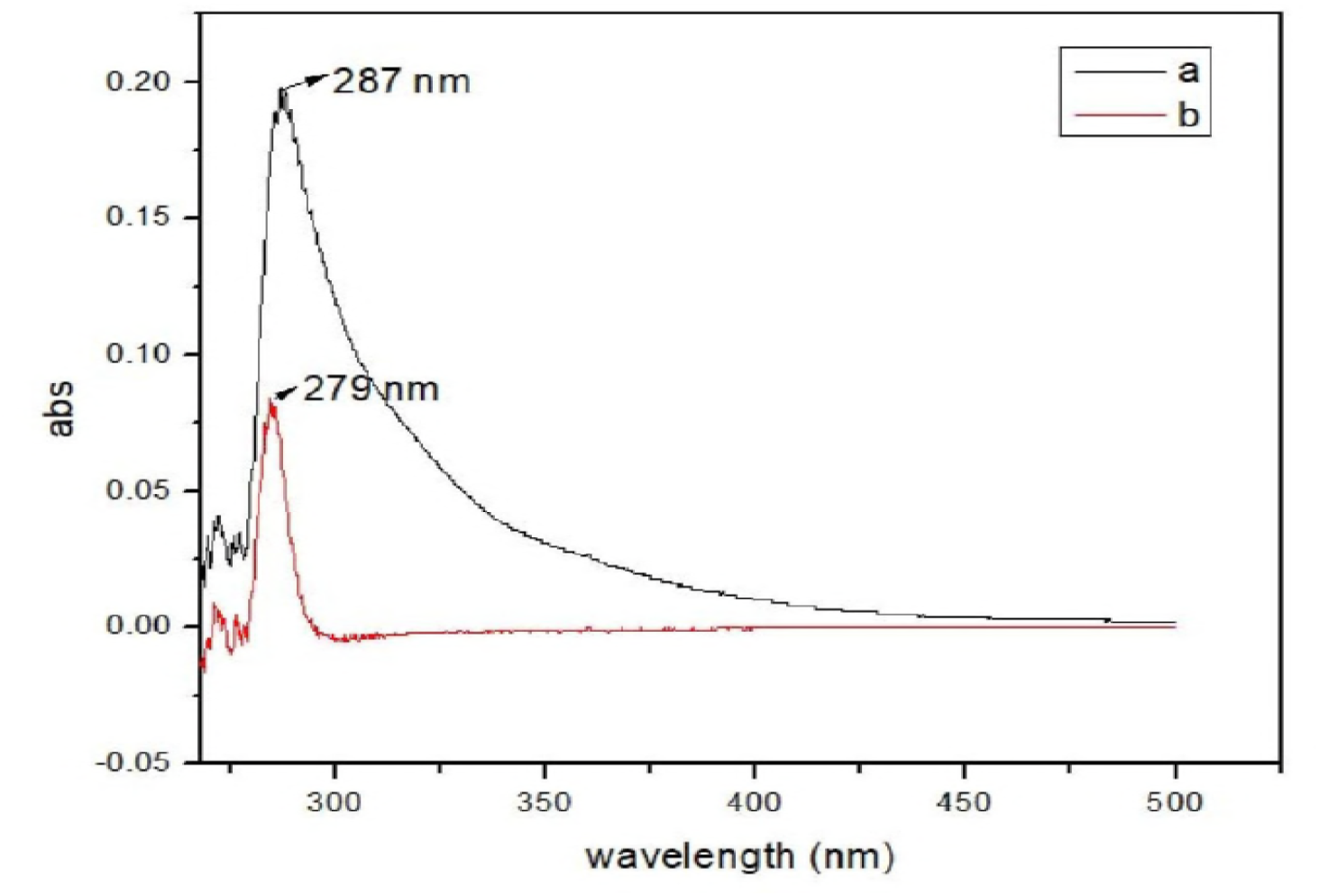
UV Spectroscopy analysis of purified bioactive compounds of F1A (a) and F1B (b) from active fraction of EtOAcE of actinomycetes CAHSH-2.

**Fig. 5b.**
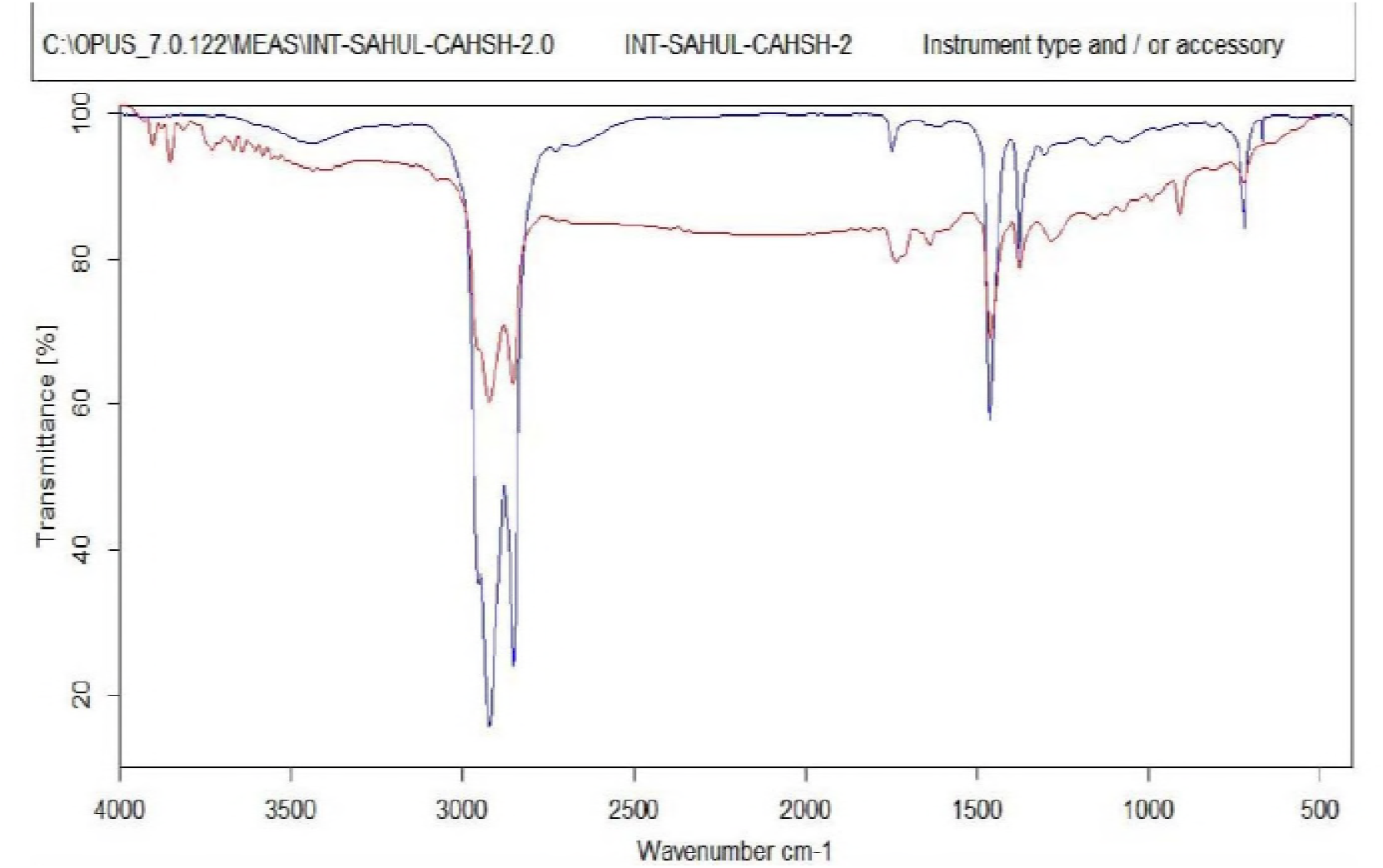
FT-IR Spectroscopy analysis of the purified bioactive compounds, F1A (Blue line) and F1B (Red line) purified from the EtOAcE of actinomycetes CAHSH-2.

**Fig. 5c.**
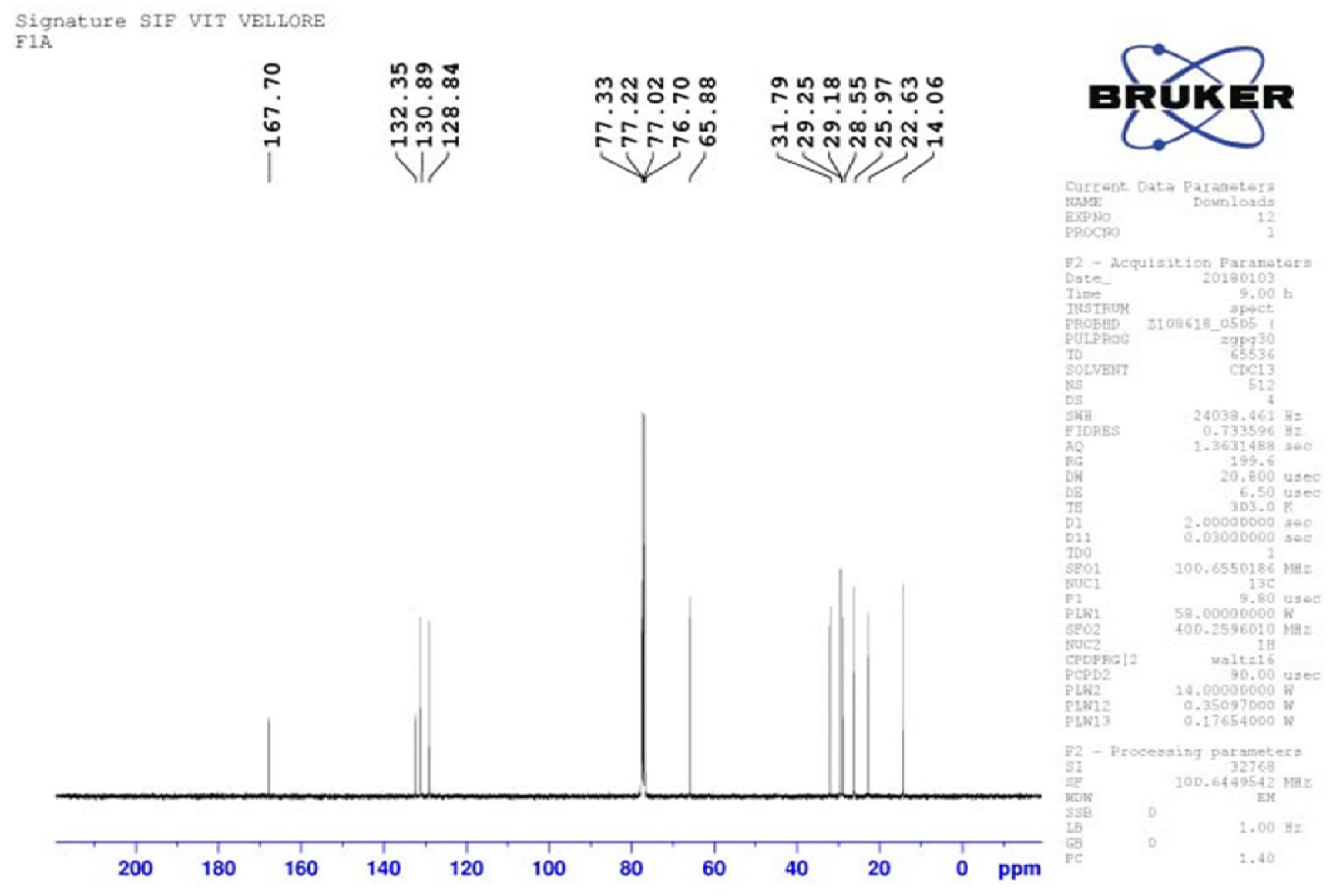
^1^H NMR spectrum analysis of the purified bioactive compound, F1A from the active fraction of EtOAcE of actinomycetes CAHSH-2.

The ^1^H NMR of the compound (F1A) showed the symmetry of the molecule. The two-terminal methyl and the two-secondary methyl at C-2^1^ appeared at δ 0.88 integrating for 12 protons. The aromatic protons H-3 and H-4 appeared at δ 7.74 and 7.73 as AA1BB1 system. The methylene protons at H-11 attached to the ester appeared at δ 4.09 as multiplet. The other methylene protons appeared at δ 1.43-1.33 (Fig 5d).

**Fig. 5d.**
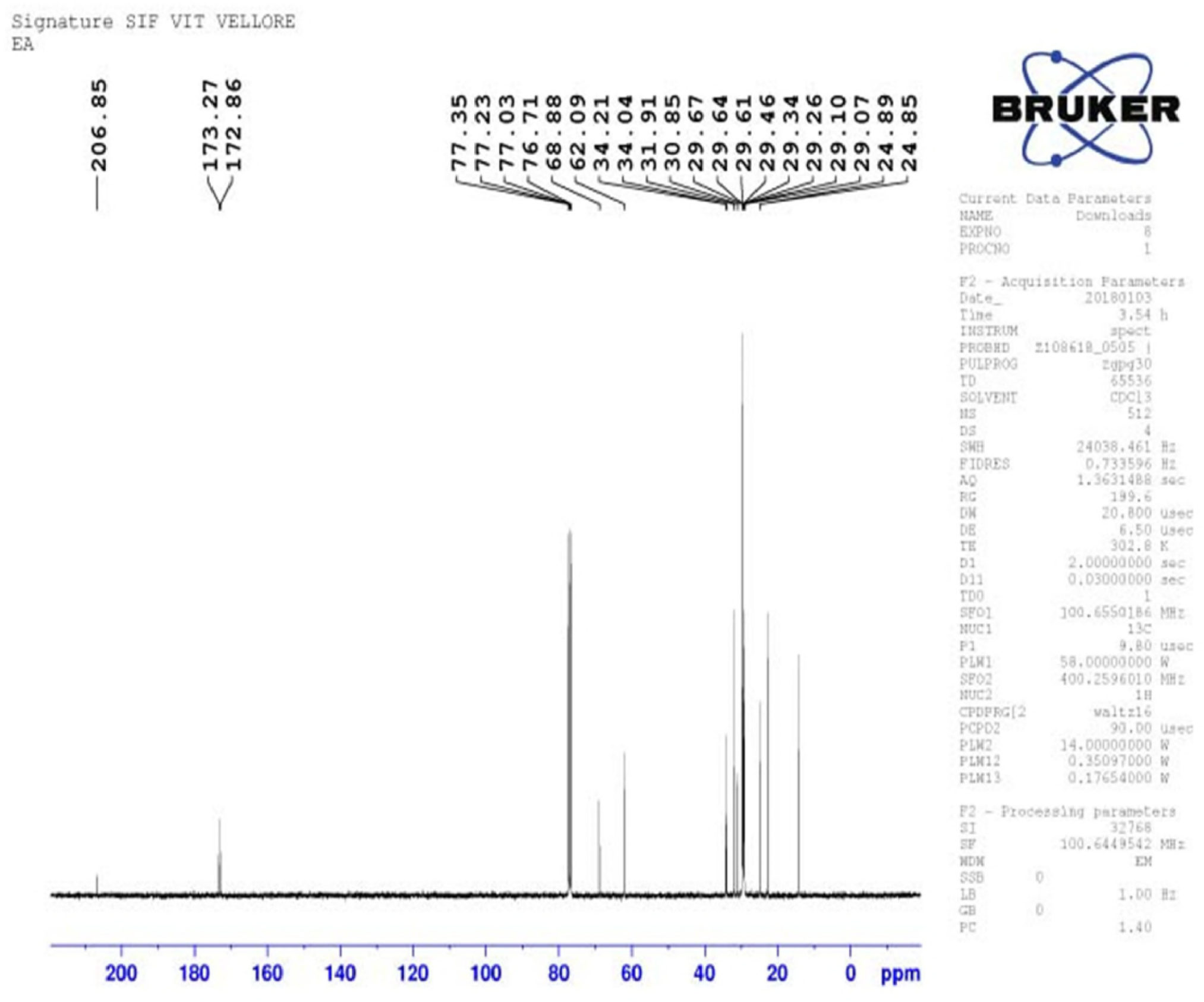
^13^C NMR spectrum analysis of the purified bioactive compound, F1A from the active fraction of EtOAcE of actinomycetes CAHSH-2.

The ^13^C NMR spectrum also confirmed the structure. The ester carbonyl appeared at δ 167.7 while the methylene attached to ester oxygen appeared δ 68.1. The terminal methyl appeared at δ 14.1 while the methyl attached to C-2^1^ appeared at δ 10.9. The three aromatic carbons C-2, C-3 and C-4 appeared at δ 130.9, 128.8 and 132.4, respectively. The other carbons are also accounted for 167.70, 132.35, 130.89, 128.84, 65.88, 31.79, 29.25, 29.18, 28.55, 25.97, 22.63, 14.06 ppm (Fig 5e). The compound having the antiviral activity against WSSV was identified as Di-N-Octyl phthalate. based on the interpretation of the results of GC-MS, FTIR and NMR spectrum analyses.

**Fig 5e.**
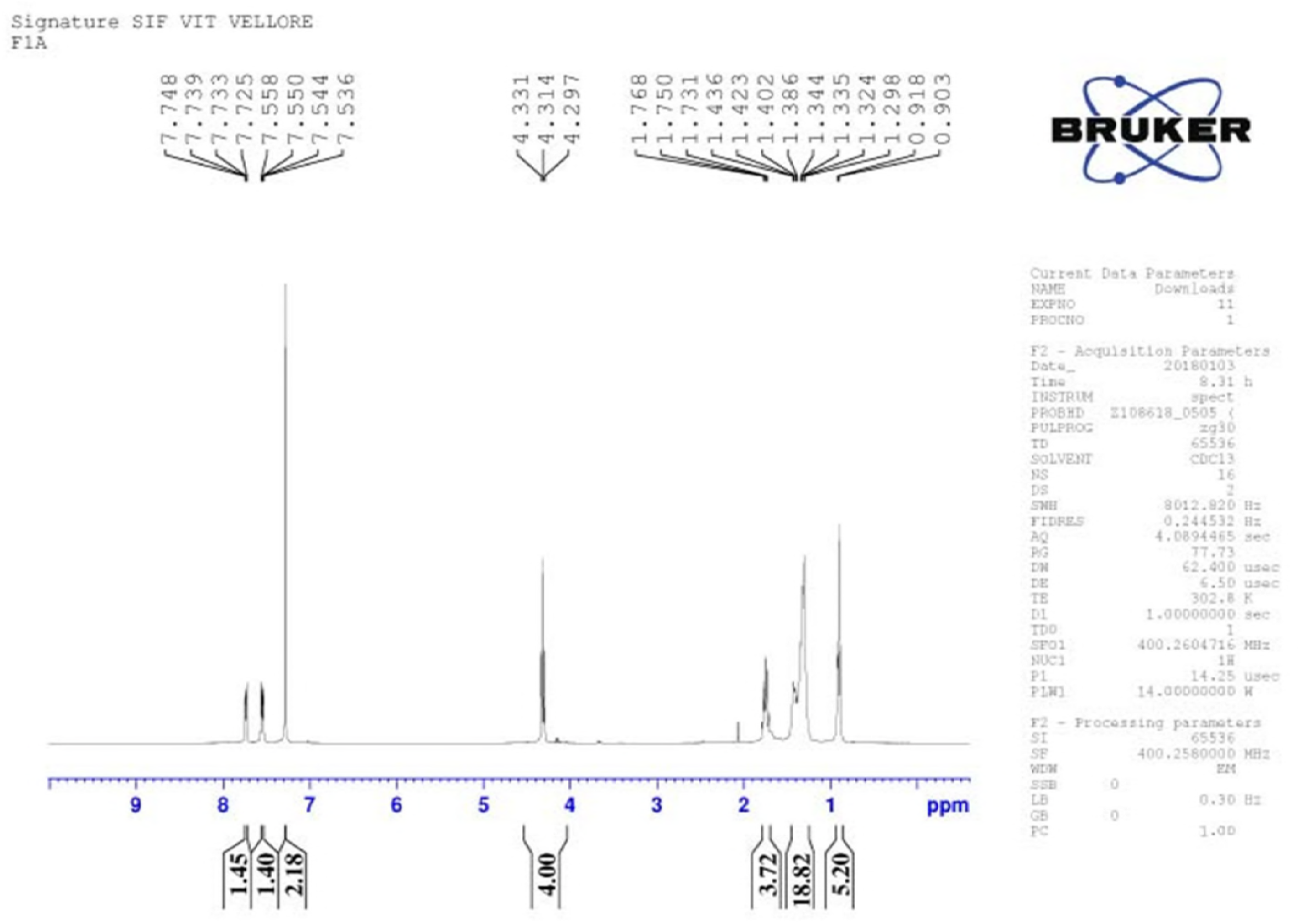
^1^H NMR spectrum analysis of the purified bioactive compound, FIB from the active fraction of EtOAcE of actinomycetes CAHSH-2.

The ^1^H NMR of the compound (F1B) showed the symmetry of the molecule. The two-terminal methyl and the two-secondary methyl at C-2^1^ appeared at δ 0.88 integrating for 12 protons. The aromatic protons H-3 and H-4 appeared at δ 7.67 and 7.52 as AA1BB1 system. The methylene protons at H-11 attached to the ester appeared at δ 4.09 as multiplet. The other methylene protons appeared at δ 1.43-1.33 (Fig 5f).

**Fig. 5f.**
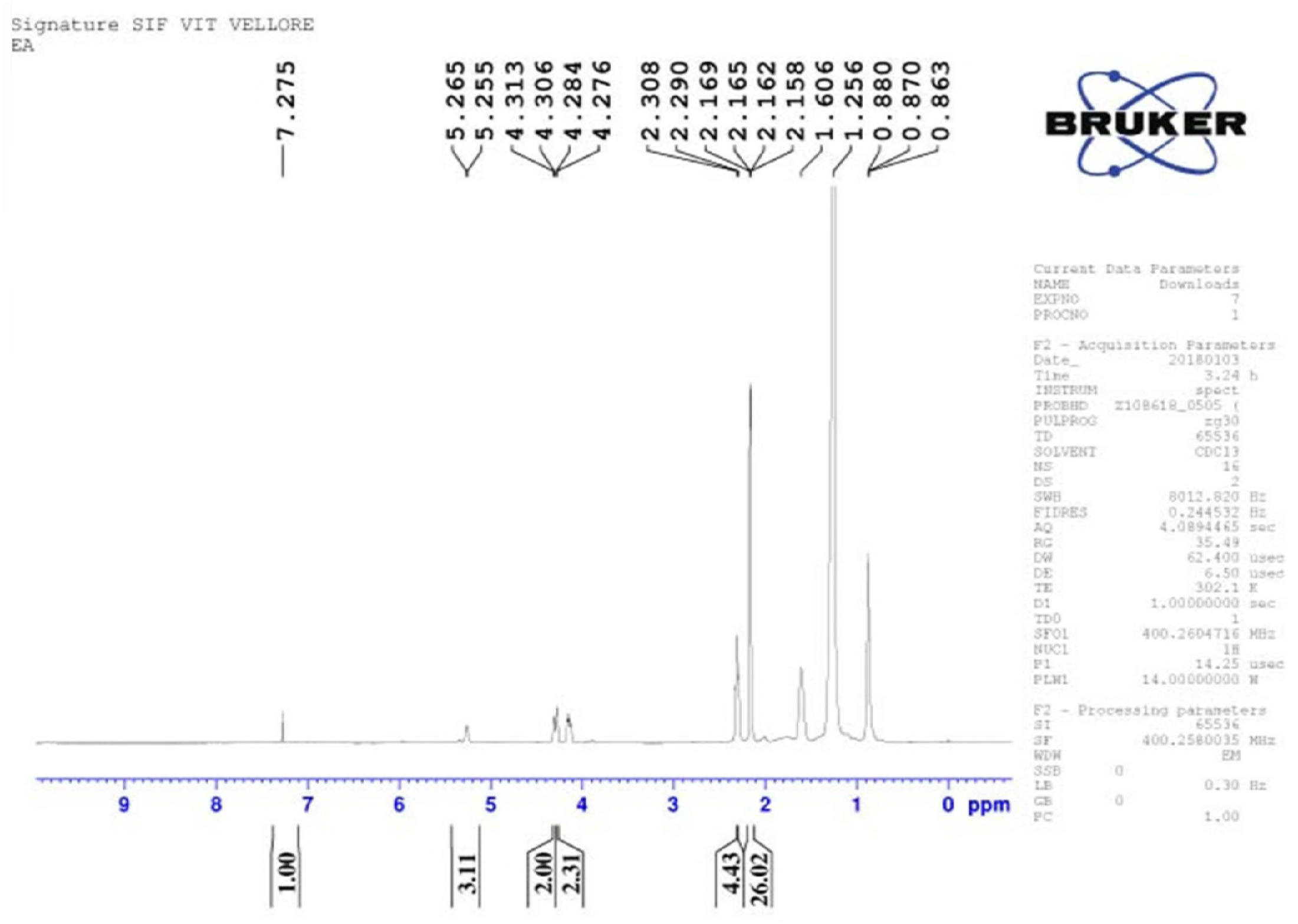
^13^C NMR spectrum analysis of the purified bioactive compound, F1B from the active fractio n of EtOAcE of actinomycetes CAHSH-2.

The ^13^C NMR spectrum also confirmed the structure. The ester carbonyl appeared at δ 167.7 while the methylene attached to ester oxygen appeared δ 68.1. The terminal methyl appeared at δ 14.1 while the methyl attached to C-2^1^ appeared at δ 10.9. The three aromatic carbons C-2, C-3 and C-4 appeared at δ 130.9, 128.8 and 132.4, respectively. The other carbons are also accounted for 167.7 (C-1), 68.1 (C-11), 38.7 (C-21), 23.7 (C-31), 28.9 (C-41), 30.3 (C-51), 22.9 (C-61), 14.1 (C-71), 10.9 (CH-CH3) (Fig 5g). The F1A having limited antiviral activity against WSSV was identified as bis (2-methylheptyl) phthalate.

**Fig. 5g.**
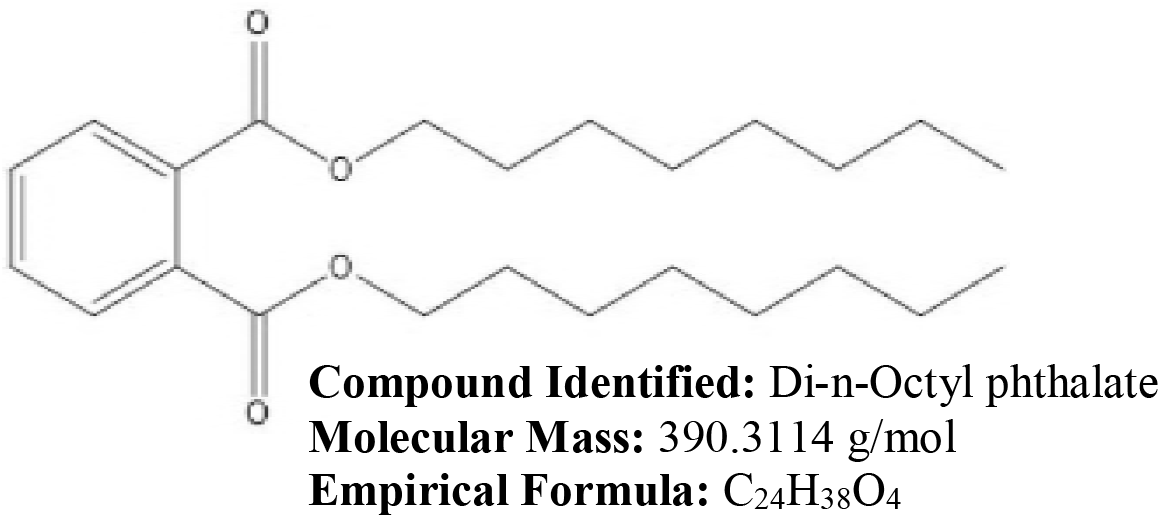
Molecular structure of the purified bioactive compound, Di-n-Octyl phthalate isolated from active fraction of EtOAcE of *Streptomyces* sp. CAHSH-2.

### 3.5. Molecular docking studies

In this study, the aim was to target VP26 and VP28 proteins of WSSV. FTsite identified three ligand binding sites for these target proteins (Fig 6a). A significant variation among these three ligand binding sites was observed. Among these three sites obtained from FTsite, site 1 was conserved in VP 26 and VP28 and therefore site 1 was selected for molecular docking study. The ligand binding sites in VP26 are TYR46, GLN48, MET49, MET50, ARG51, ARG67, TYR69, ASN70, THR71, PRO72 and ligand binding sites in VP28 are GLU50, ASN51, LEU52, ARG53, PRO55, VAL70, PHE72, ASP73, SER74, ASP75, THR76 and ILE82. The 3D structure of probable ligand binding site is shown in surface model and amino acid residues located at ligand binding site of VP 26 and VP28 are shown in (Fig 6b). The optimized di-n-octyl phthalate and bis (2-methylheptyl) phthalate structures were docked into the ligand binding site of VP26 and VP28 (Fig 6c). The docking results were analysed and results revealed that the lowest binding energy value represents stabled protein-ligand complex compared to higher binding energy value. Table 4, reveals that a variation was observed in the binding affinity of di-n-octyl phthalate and bis (2-methylheptyl) phthalate with VP26 and VP28 viral proteins. The di-n-octyl phthalate showed the lowest binding energy values with VP26 and VP28 (−5.54 Kcal/Mol and −4.7 Kcal/Mol) when compared to bis (2-methylheptyl) phthalate. The di-n-octyl phthalate was found to have high binding affinity with VP26 and VP28, whereas binding affinity of the bis (2-methylheptyl) phthalate with VP26 and VP28 was low. The binding poses of di-n-octyl phthalate and bis (2-methylheptyl) phthalate into VP26 and VP28 proteins of WSSV are shown in (Fig 6d–g)

**Fig. 6a.**
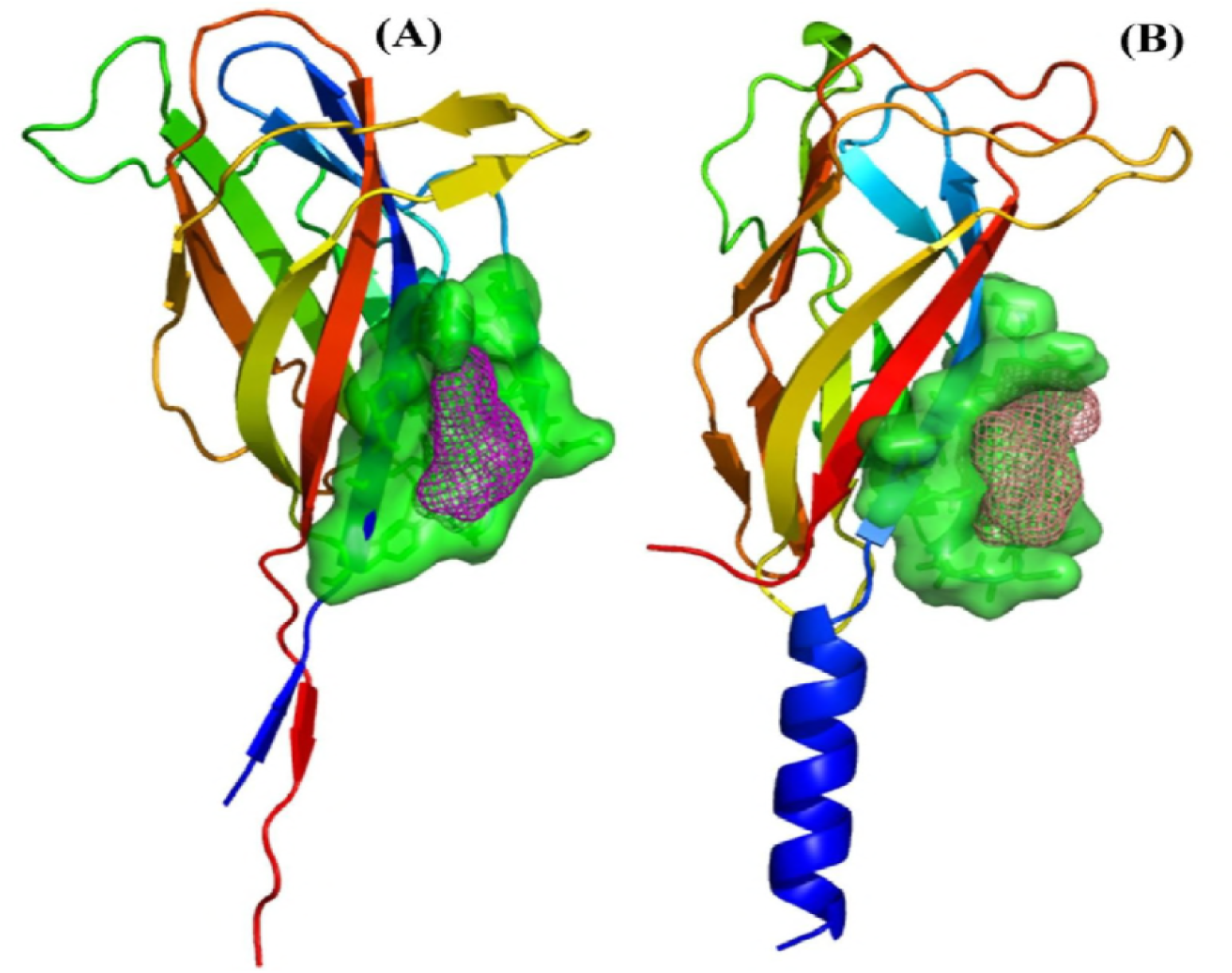
3D Structure of VP26 (A) and VP28 (B). Ligand binding site is obtained by FTSite and active site residues are shown in green colour surface model. Ligand binding pocket are show in mesh model.

**Fig. 6b.**
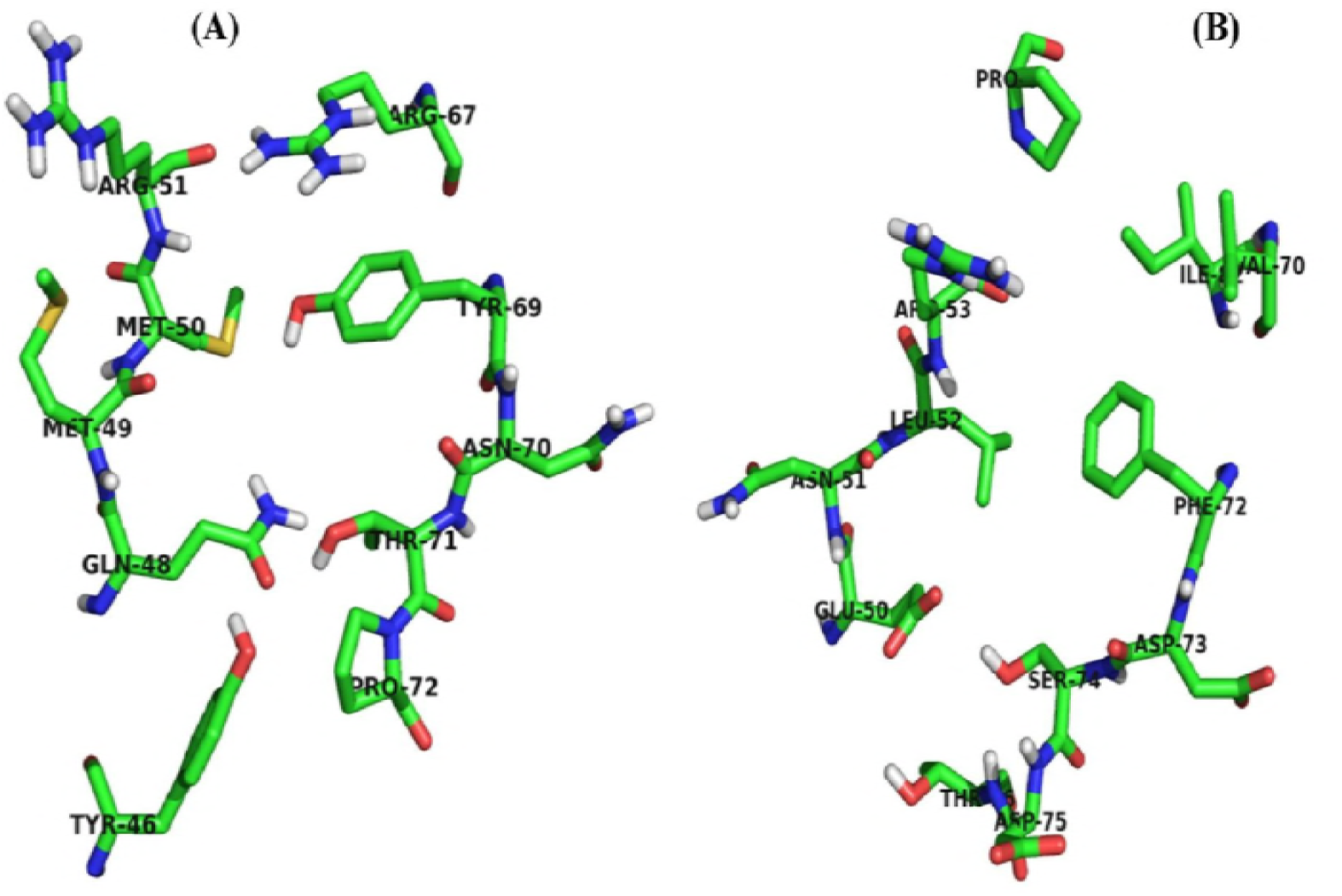
Amino acid residues located at active site of VP26 (A) and VP28 (B).

**Fig. 6c.**
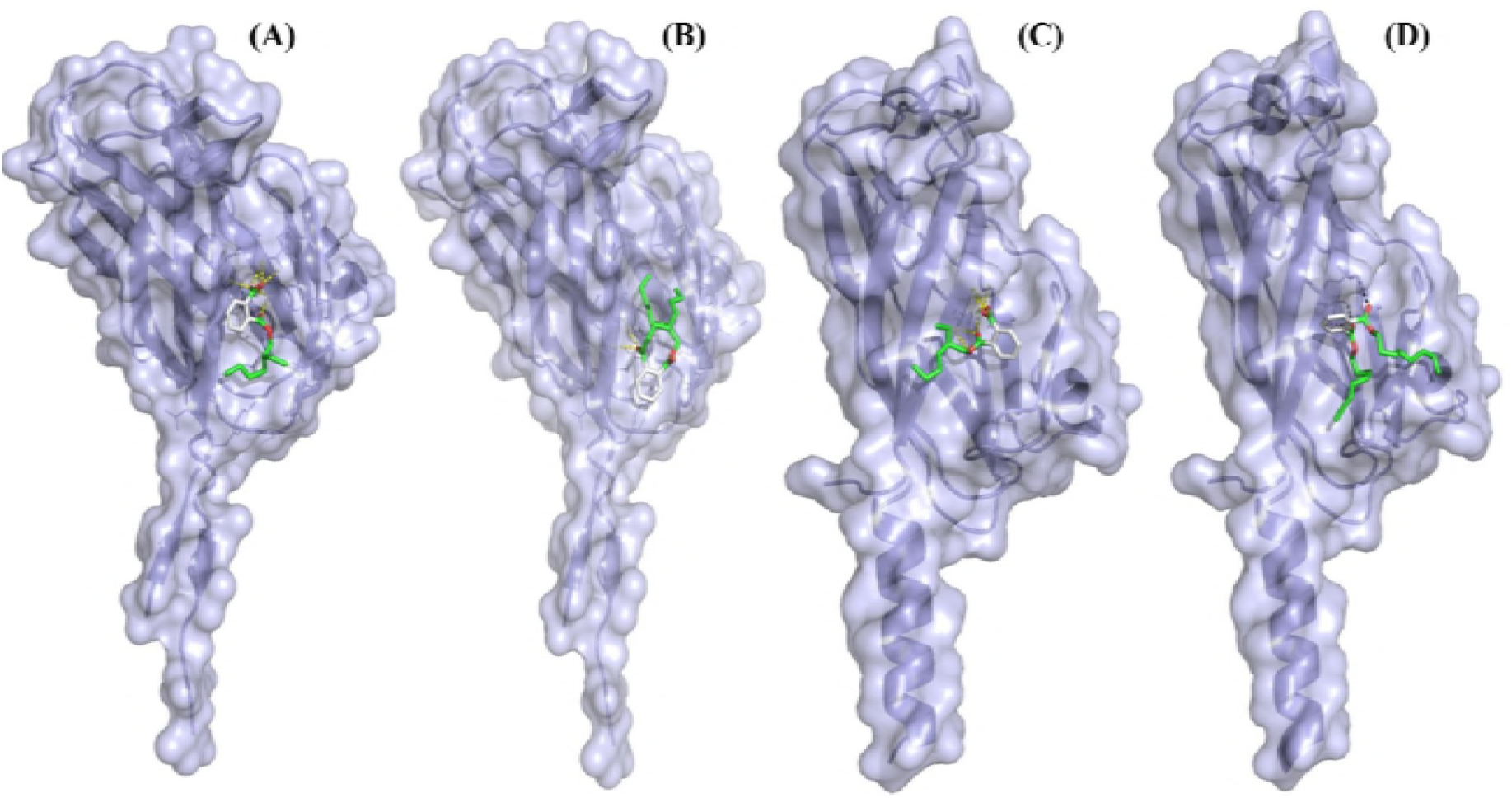
Molecular docking of di-n-octyl phthalate and bis(2-methylheptyl) phthalate with VP26 and VP28. Binding pattern of di-n-octyl phthalate with VP26 (A), Binding pattern of bis (2-methylheptyl) phthalate with VP26 (B), Binding pattern of di-n-octyl phthalate with VP28 (C) and Binding pattern of bis (2-methylheptyl) phthalate with VP28 (D).

**Fig. 6d.**
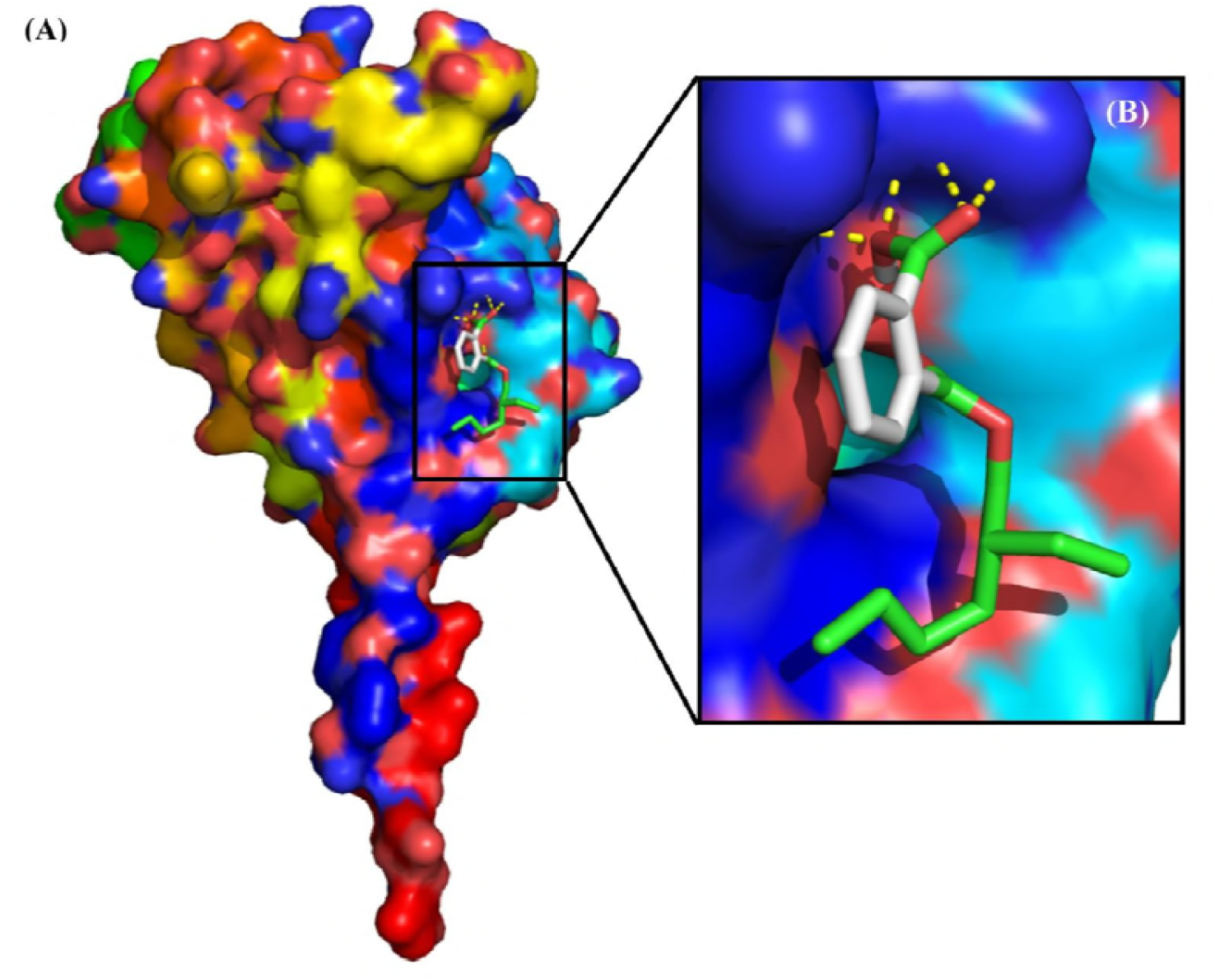
Binding pose of di-n-octyl phthalate in the binding site of VP26 (A). A close-up view of the binding pose of di-n-octyl phthalate (B).

**Fig. 6e.**
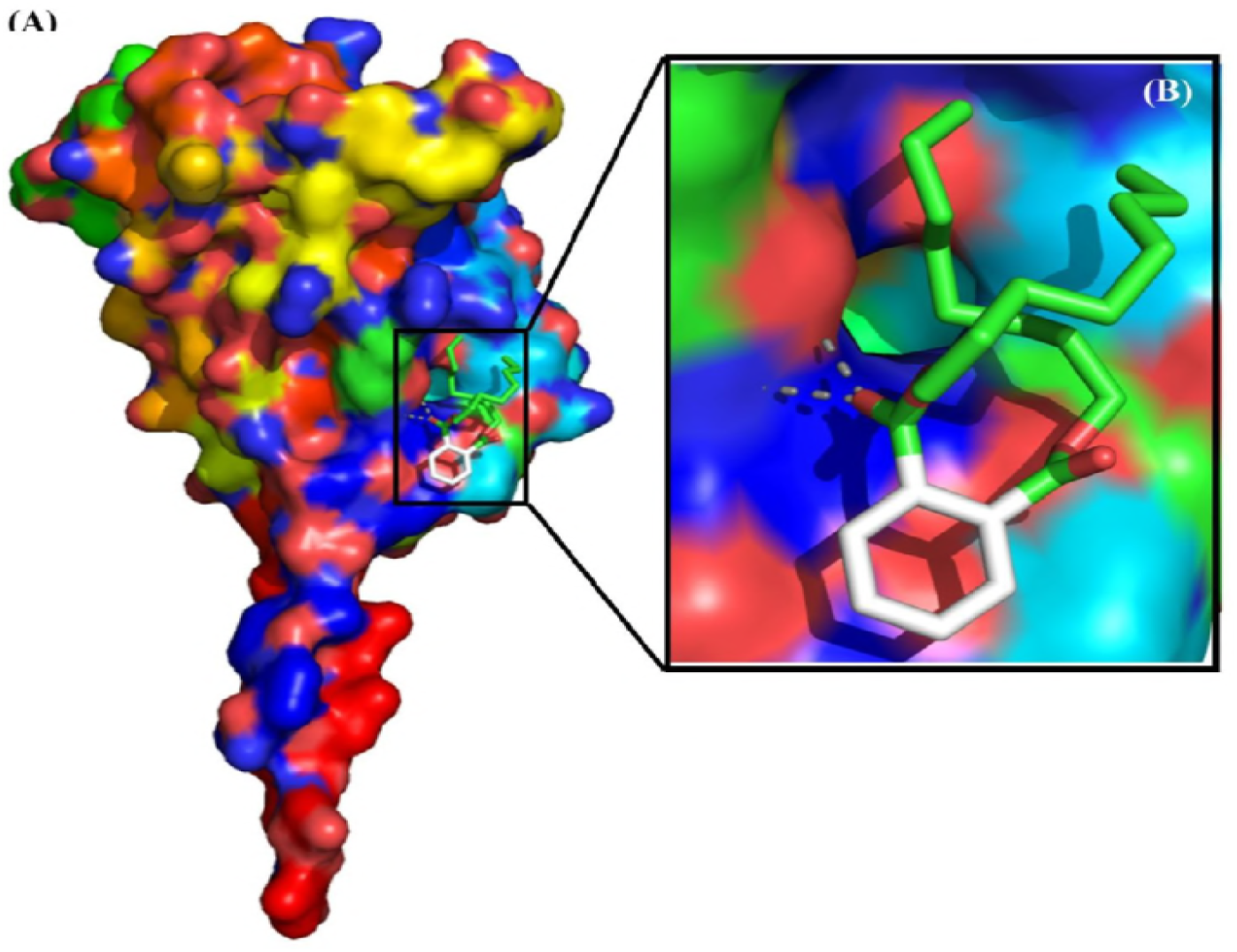
Binding pose of bis (2-methylheptyl) phthalate in the binding site of VP26 (A) and A close-up view of the binding pose of bis (2-methylheptyl) phthalate (B).

**Fig. 6f.**
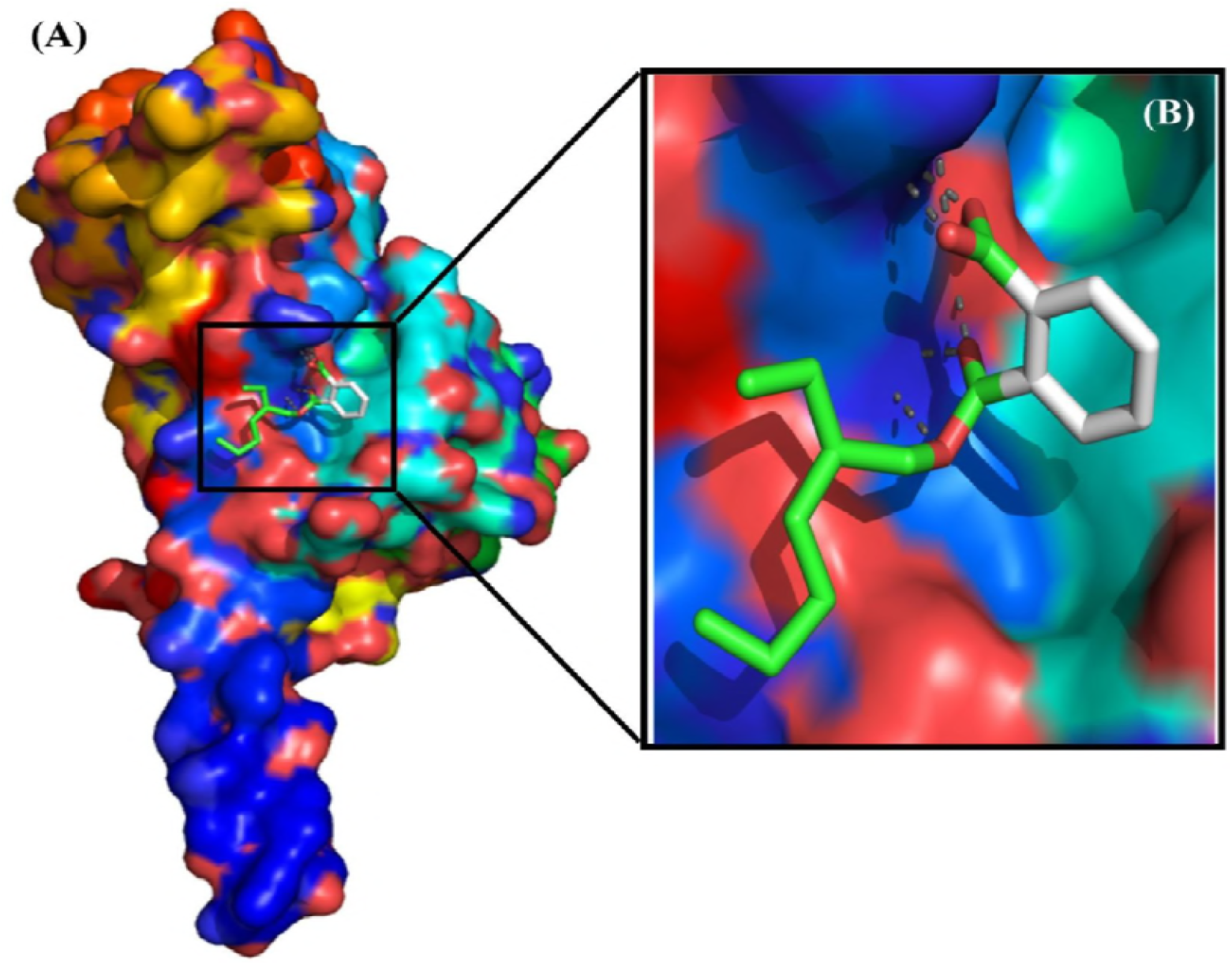
Binding pose of di-n-octyl phthalate in the binding site of VP28 (A) and a close-up view of the binding pose of di-n-octyl phthalate (B).

**Fig. 6g.**
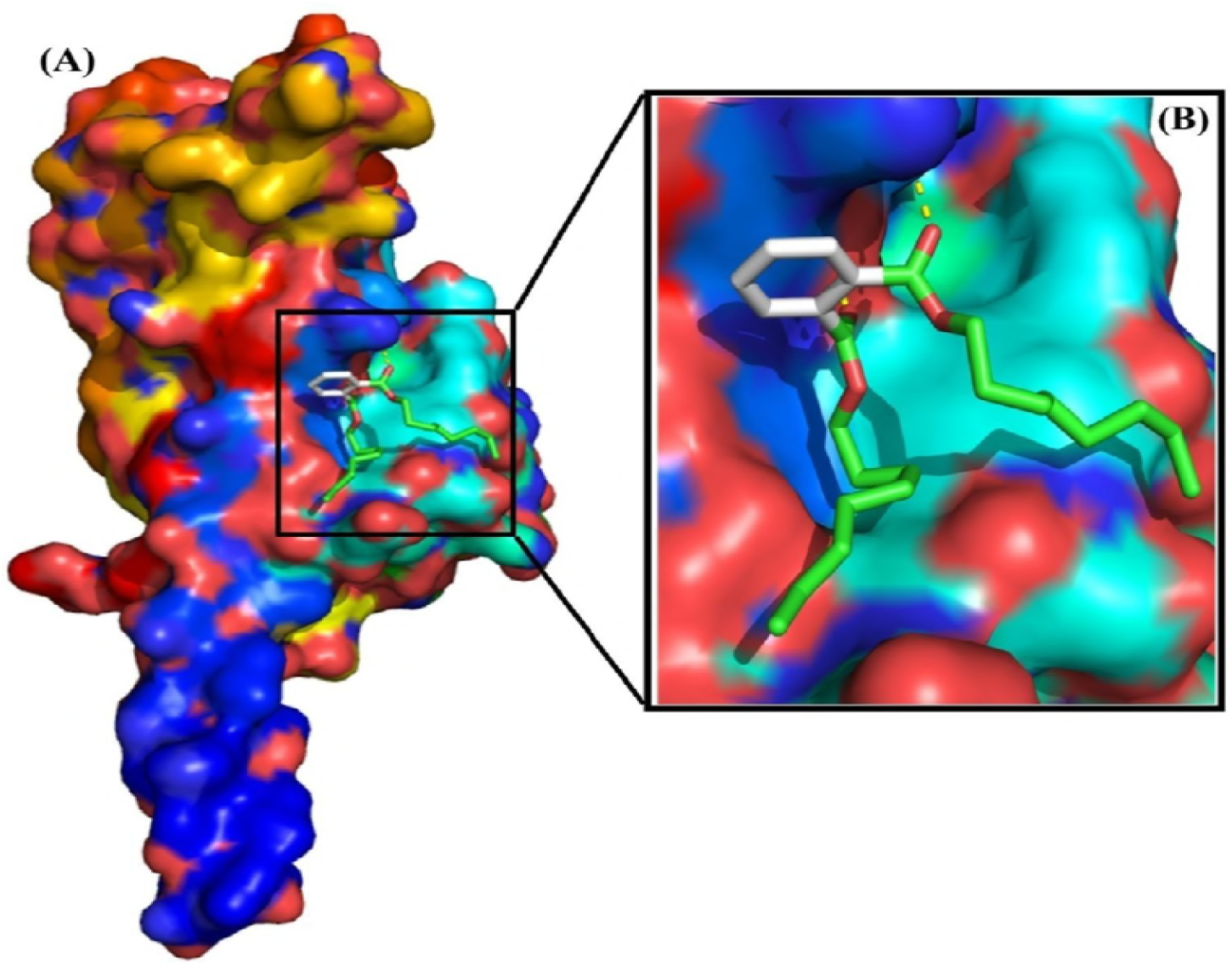
Binding pose of bis (2-methylheptyl) phthalate in the binding site of VP28 (A) and a close-up view of the binding pose of bis (2-methylheptyl) phthalate (B).

**Table 4.** Auto dock results of di-n-octyl phthalate and bis(2-methylhepty) phthalate against VP26 and VP28

| Parameters | di-n-octyl phthalate - VP26 | bis(2-methylhepty) phthalate VP26 | di-n-octyl phthalate - VP28 | bis(2-methylhepty) phthalate VP28 |
| --- | --- | --- | --- | --- |
| Binding energy Kcal/Mol | -5.54 | -3.57 | -4.7 | -3.72 |
| Ligand efficiency | -0.28 | -0.13 | -0.24 | -0.13 |
| Inhibit constant | 87.47 | 2.43 | 360.06 | 1.87 |
| Intermol_energy | -7.31 | -5.92 | -6.34 | -6.93 |
| Vdw_hb_desolv_energy | -5.73 | -5.86 | -5.65 | -6.83 |
| Electrostatic energy | -1.57 | -0.07 | -0.69 | -0.1 |

Further investigation was carried out on docked complexes to check H-bond network between ligands and viral proteins. From the interaction analysis, a difference was noticed in the formation of H-bond between the compounds [di-n-octyl phthalate and bis (2-methylheptyl) phthalate] and viral proteins (VP26 and VP28) (Fig 6h [A-D]). Upon analysis of the interaction of di-n-octyl phthalate with VP26, it was observed that the amino acid residues Arg67, Tyr69 and Arg51 contributed four H-bonds with di-n-octyl phthalate whereas in VP28, the ligand binding site residues Met49 and Thr71 participate in the H-bond interaction. On analyzing di-n-octyl phthalate - VP28 complex, it was seen that only one cationic amino acid residue Arg53 was involved in the formation of seven H-bond interactions with di-n-octyl phthalate. The interaction analysis of bis (2-methylheptyl) phthalate - VP28 complex reveals that the cationic residue Arg53 contributed three H-bonds with bis (2-methylheptyl) phthalate. Binding pose of di-n-octyl phthalate and bis (2-methylheptyl) phthalate with VP26 and VP28 viral proteins and close up view of each complex is shown in surface model. In the present study, on analyzing VP26 and VP28 in complex with di-n-octyl phthalate and bis (2-methylheptyl) phthalate, di-n-octyl phthalate showed higher binding affinity and multiple hydrogen bonds with VP26 and VP28 viral proteins.

**Fig. 6h.**
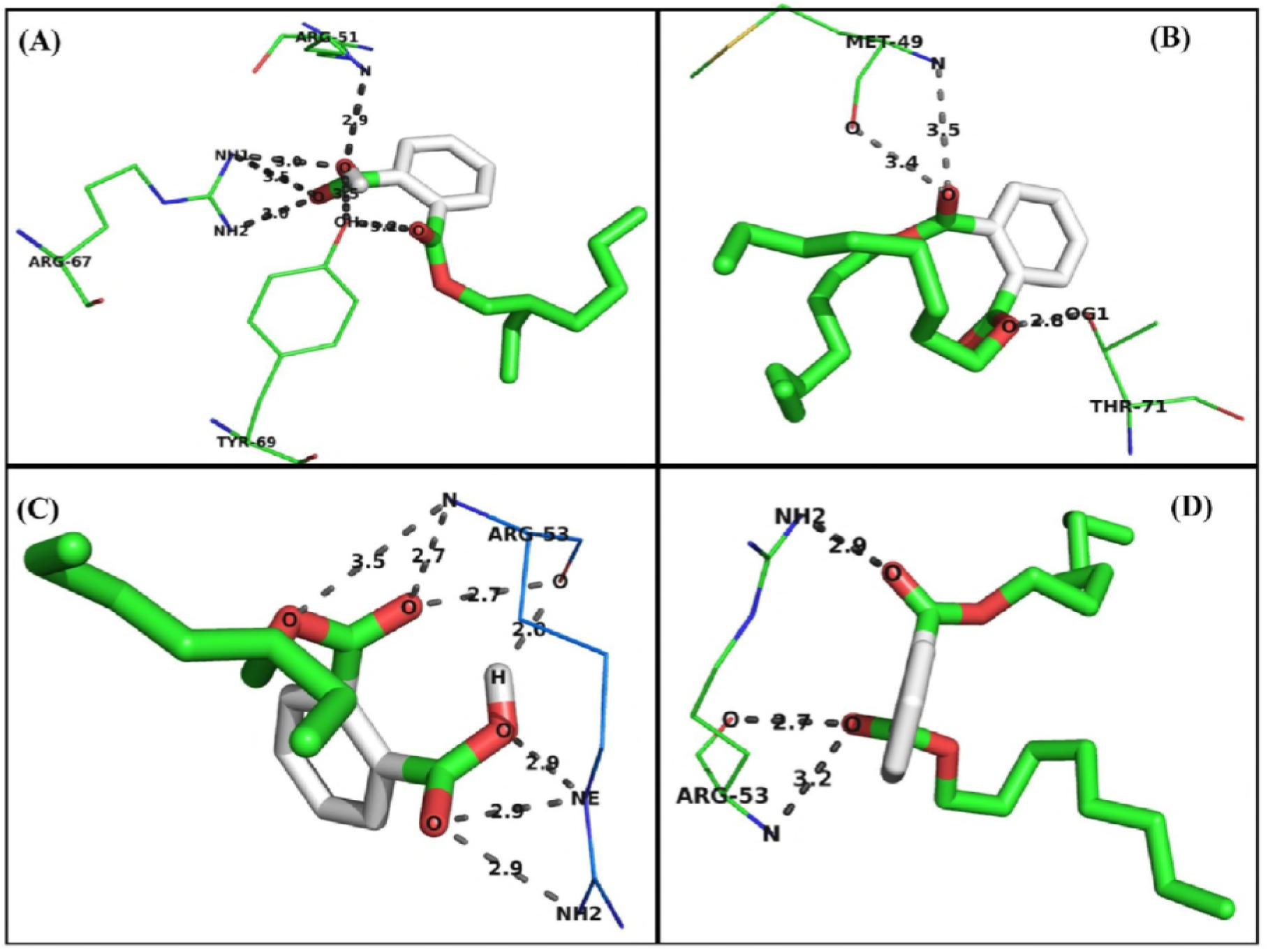
Molecular interaction of di-n-octyl phthalate and bis (2-methylheptyl) phthalate with VP26 and VP28. (A). Interaction of di-n-octyl phthalate with VP26. (B) Interaction of bis (2-methylheptyl) phthalate with VP26. (C) Interaction of di-n-octyl phthalate with VP28. (D) H-bond network between bis (2-methylheptyl) phthalate and VP28.

### 3.6. Confirmation of antiviral activity of potential actinomycetes isolates

The PCR results showed WSSV-positive in shrimp injected with EtOAcE of CAHSH-2 treated WSSV till 10 d p.i. and became negative on 15 d p.i (Fig 7a). The RT-PCR analysis revealed that the expression of VP28 gene of WSSV was gradually decreased and totally disappeared after 5 d p.i. in shrimp injected CAHSH-2 treated WSSV (Fig 7b). The shrimp injected with EtOAcE of CAHSH-2 treated WSSV showed negative to WSSV after 5 d p.i. by Western blot (Fig 7c–d). In ELISA, the OD value corresponding to the presence of WSSV (VP28 protein) in shrimp injected with WSSV treated with EtOAcE of CAHSH-2 was found to decrease gradually and became negligible after 5 d p.i (Fig 7e).

**Fig. 7a.**
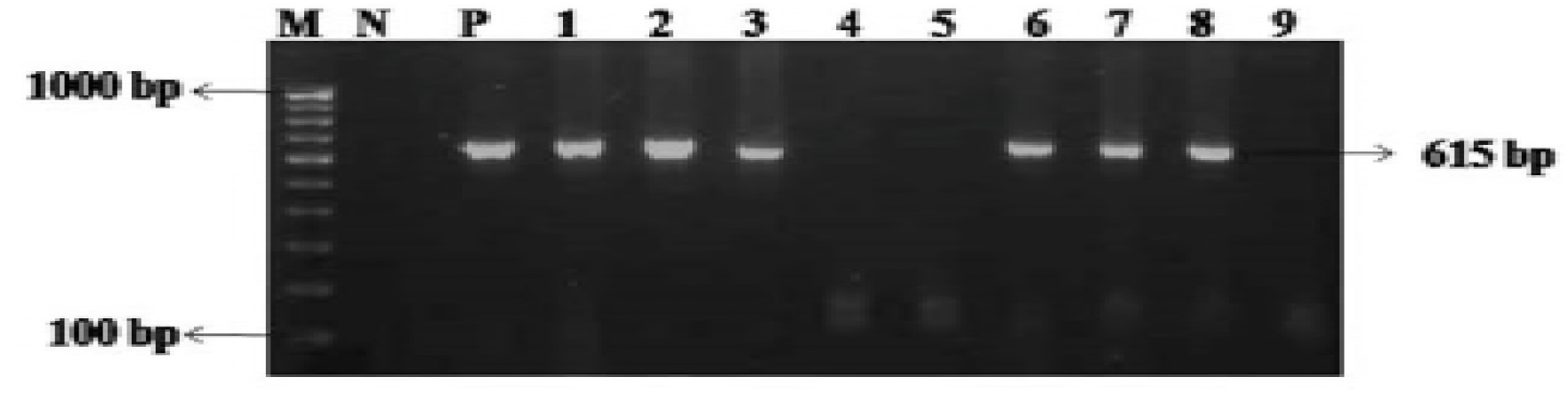
PCR detection of WSSV in the gill and head tissues of *L. vannamei* injected with EtOAcE of CAHSH-2 treated WSSV at different time intervals. Lane M-DNA 100 bp Marker, Lane N - Negative Control, Lane P – Positive Control, Lane 1 – 3^rd^ day gill, Lane 2–5^th^ day gill, Lane 3–10^th^ day gill, Lane 4–15^th^ day gill, Lane 5 – Negative control, Lane 6 – 3^rd^ day head tissue, Lane 7–5^th^ day head tissue, Lane 8 – 10^th^ day head tissue, Lane 9–15^th^ day head tissue.

**Fig. 7b.**
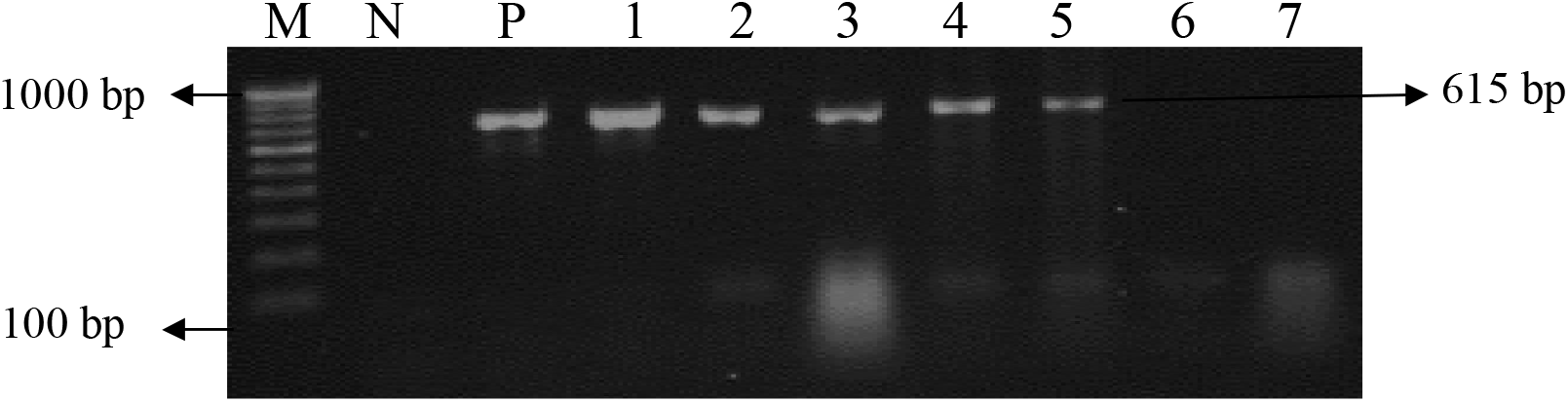
RT-PCR detection of VP28 transcript in the gill tissue of *L. vannamei* injected with EtOAcE of CAHSH-2 treated WSSV at different time intervals. Lane M–DNA 100 bp Marker, Lane N – Negative Control, Lane P – Positive Control, Lane 1–1^st^ day, Lane 2 – 2^nd^ day, Lane 3–3^rd^ day, Lane 4–5^th^ day, Lane 5–10^th^ day, Lane 6–13^th^ day, Lane7–15 ^th^ day.

**Fig. 7c.**
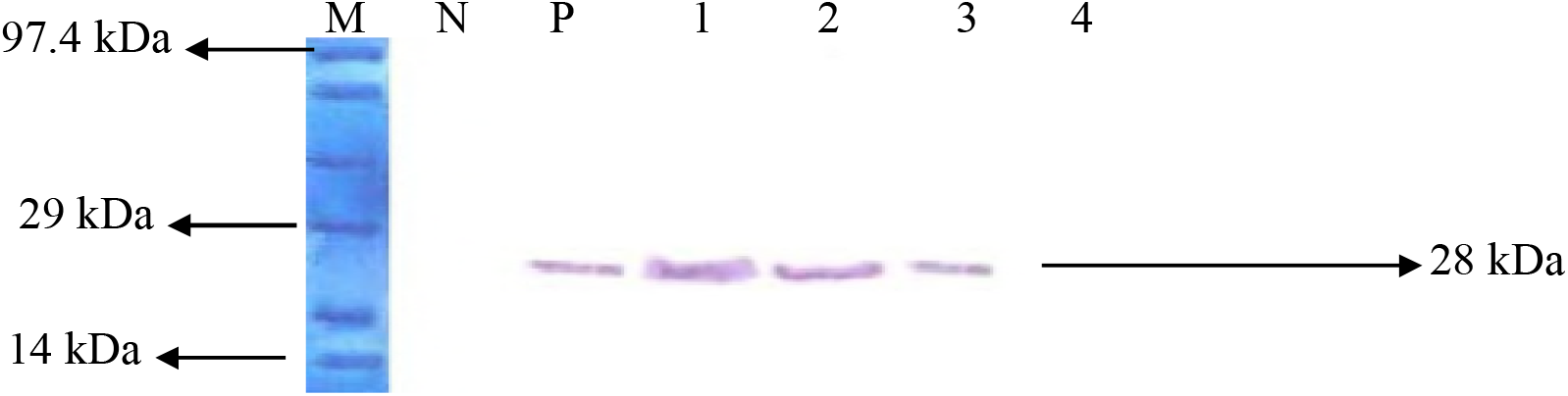
Detection of WSSV by Western blot analysis using anti-rVP28 of WSSV in the gill tissue of *L. vannamei* injected with EtOAcE of CAHSH-2 treated WSSV at different time intervals. Lane M – Protein marker; Lane N – negative control; Lane P – positive control; Lane 1 – 2^nd^ day, Lane 2 – 5 ^th^ day, Lane 3–10^th^ day, Lane 4–15^th^ day.

**Fig. 7d.**
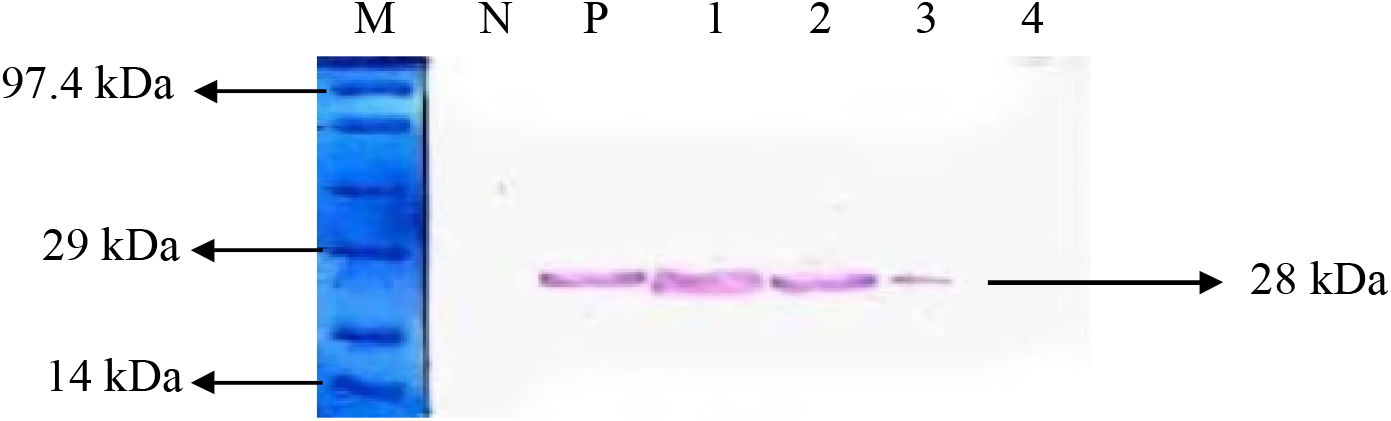
Detection of WSSV by Western blot analysis using anti-rVP28 of WSSV in the head tissue of *L. vannamei* injected with EtOAcE of CAHSH-3 treated WSSV at different time intervals. Lane M – Protein marker; Lane N – negative control; Lane P – positive control; Lane 1 – 2^nd^ day, Lane 2 – 5 ^th^ day, Lane 3 – 10^th^ day, Lane 4–15^th^ day

**Fig. 7e.**
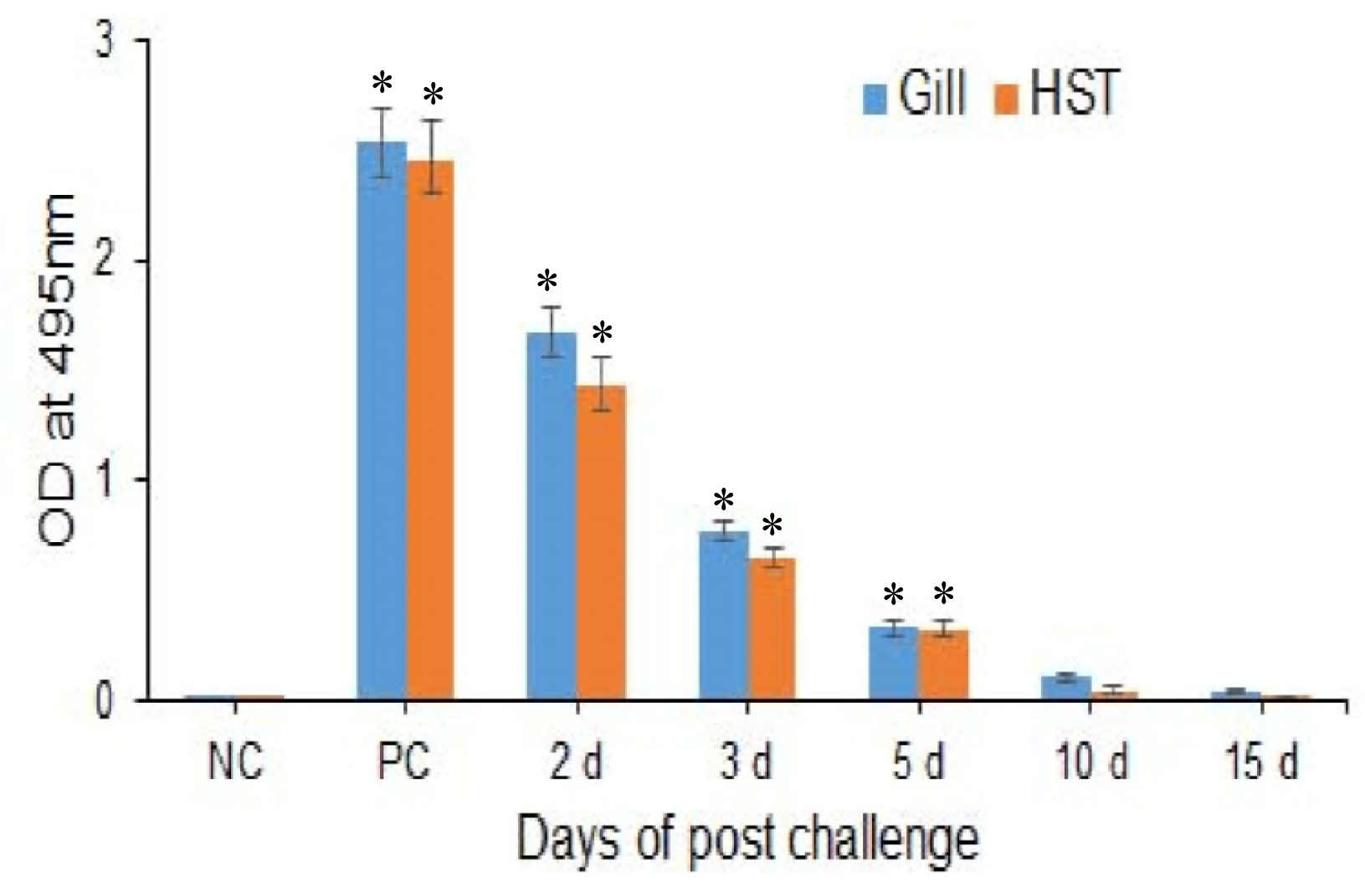
Detection of WSSV by ELISA using anti–rVP28 of WSSV in the gill and head tissues of *L. vannamei* injected with EtOAcE of CAHSH-2 treated WSSV at different time intervals.

The load of WSSV was quantified in gill and head tissues of shrimp injected with EtOAcE of CAHSH-2 treated WSSV by real time PCR at different time intervals. In the gill tissue, the WSSV load was estimated to be about 2.7 × 10^5^, 5.4 × 10^3^, 150 and 0.97 copies per mg of tissue at 2, 5, 10 and 15 d p.i., respectively. In the head tissue, the WSSV load was estimated to be about 3.1 × 10^4^, 2.7 × 10^3^, 19 and 0.96 copies per mg of tissue at 2, 5, 10 and 15 d p.i., respectively. In the shrimp injected with WSSV, the WSSV was estimated to be about 6.2 × 10^9^ in gill tissue 4.3 × 10^7^ in head tissue at moribund stage (48 h p.i.) (Table 5).

**Table 5.**
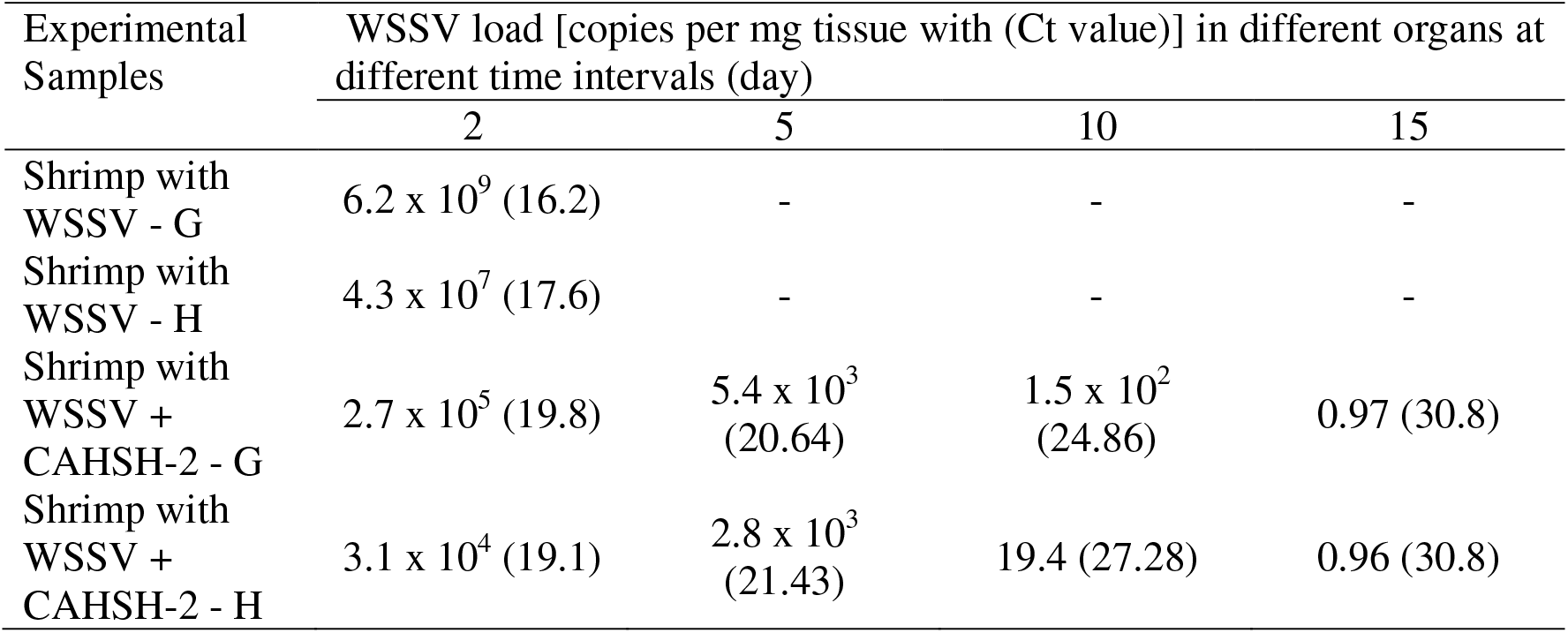
WSSV load in different organs of shrimp injected with EtOAcE of CAHSH-2 treated WSSV by quantitative real-time PCR at different time intervals

### 3.7. Toxicity studies

The toxicity of the ethyl acetate extract of actinomycetes isolate (CAHSH-2) and antiviral compound (Di-N-Octyl phthalate) purified from CAHSH-2 isolate was studied *in vivo* using *Artemia* nauplii, and shrimp post-larvae by immersion method and sub-adult and adult shrimp by intramuscular injection method. The EtOAcE of CAHSH-2 and pure compound purified from the extract of CAHSH-2 isolate were found to be non-toxic to the *Artemia* nauplii, post-larvae and adult *L. vannamei* (Table 6a). The mortality of experimental animals was found to be insignificant and similar to the negative control group of animals. In *Artemia* nauplii treated with EtOAcE of CAHSH-2 at the concentration of 10 mg or 50 mg per litre, the survival percentage ranged from 97.3 to 100% while the survival ranged from 98.73 to 99.46% in *Artemia* treated with pure compound during 48 hours of exposure. In shrimp post-larvae, the survival percentage ranged from 94.67 to 100 in the case of EtOAcE-CAHSH-2 treatment at the concentration of 10 mg or 50 mg per litre and 98.73 to 99.46 in the case of pure compound treatment at the concentration of 1 or 2 mg per litre for 48 hours of exposure (Table 6b). The toxicity of extract and pure compound was also tested in adult *L. vannamei* by intramuscular injection. The survival percent was found to be 96.67 and 93.33 in shrimp injected with EtOAcE-CAHSH-2 at the concentration of 1 mg/shrimp and 2 mg/shrimp, respectively after 10 days of treatment. In the case of pure compound, the percent survival was found to be 96.67 in shrimp injected with 2 mg of pure compound per shrimp after 10 days of treatment.

**Table 6.a.**
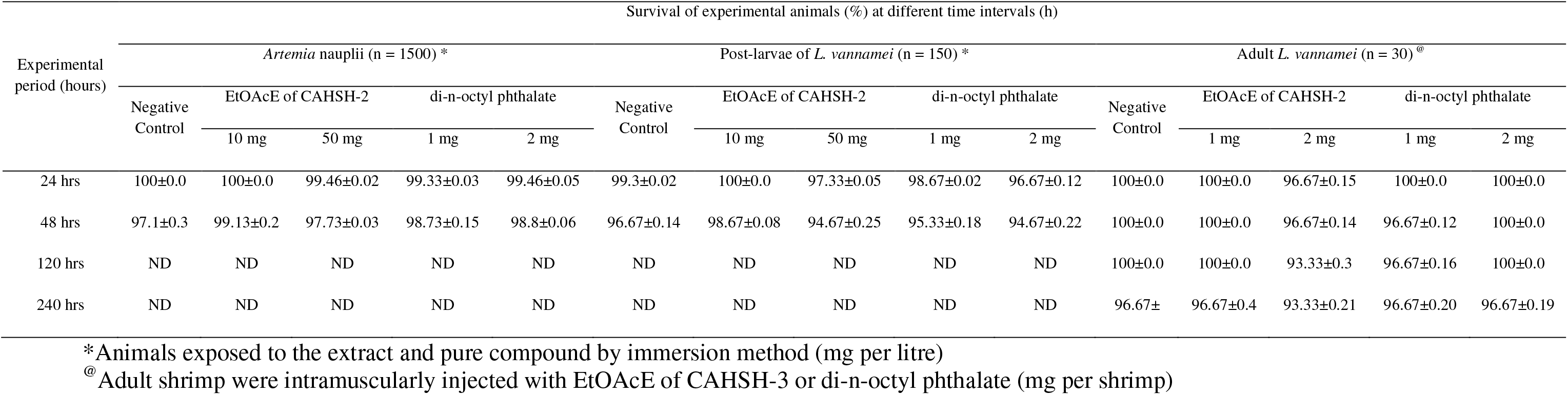
Toxicity of ethyl acetate extract (EtOAcE) of actinomycetes isolate (CAHSH-2) and di-n-octyl phthalate having antiviral activity against WSSV tested in *Artemia* nauplii, post-larvae and adult of *Litopenaeus vannamei*.

**Table 6.b.** Toxicity of ethyl acetate extract (EtOAcE) of actinomycetes isolate (CAHSH-3) having antiviral activity against WSSV tested in *Artemia* nauplii, post-larvae and adult of *Litopenaeus vannamei*

| Experimental period (hours) | Survival of experimental animals (%) at different time intervals (h) |  |  |  |  |  |  |  |  |
| --- | --- | --- | --- | --- | --- | --- | --- | --- | --- |
|  | *Artemia* nauplii (n = 1500)* |  |  | Post-larvae of *L. vannamei* (n = 150)* |  |  | Adult *L. vannamei* (n = 30)<sup>@</sup> |  |  |
|  | Negative Control | EtOAcE |  | Negative Control | EtOAcE |  | Negative Control | EtOAcE |  |
|  |  | 10 mg | 50 mg |  | 10 mg | 50 mg |  | 1 mg | 2 mg |
| 24 hrs | 99.67±0.32 | 99.73±0.22 | 99.27±0.25 | 100±0.0 | 99.13±0.18 | 98.47±0.29 | 100±0.0 | 96.67±0.44 | 100±0.0 |
| 48 hrs | 98.47±0.58 | 98.53±0.02 | 97.67±0.37 | 98.93±0.24 | 97.73±0.11 | 97.4±0.31 | 96.67±0.42 | 96.67±0.58 | 96.67±0.36 |
| 120 hrs | ND | ND | ND | ND | ND | ND | 96.67±0.53 | 93.33±0.61 | 93.33±0.03 |
| 240 hrs | ND | ND | ND | ND | ND | ND | 93.33±3.33 | 93.33±0.74 | 86.67±0.92 |

\*Animals exposed to the extract by immersion method (mg per litre)
<sup>@</sup>Adult shrimp were intramuscularly injected with EtOAcE of CAHSH-3 (mg per shrimp)

## 4. Discussion

Disease outbreaks caused by viral and bacterial pathogens have been recognized as a major constraint to aquaculture production, trade and are responsible for severe economic loss in aquaculture industry worldwide. Various strategies (efficient diagnostics, good nutrition, efficient vaccines, immunostimulants, non-specific immune-enhancers and probiotics) have been followed to prevent or control diseases of aquatic animals in the culture systems. In spite of these strategies, some of the diseases (WSS of shrimp and EUS in fish) have been persisting for more than a decade. No control strategies have so far been developed to prevent these diseases. Hence, novel strategies need to be developed to control or prevent WSSV of shrimp at field level since it has tremendous value in shrimp culture industry worldwide. In the present study, an attempt was made to screen actinomycetes isolates for possible antiviral compounds especially for WSSV.

Several reports revealed that the actinomycetes from different habitats were found to be a good source of antiviral and antibacterial compounds against viruses and bacteria of plants, animals and humans (43, 44, 45, 46, 47, 48, 49, 50, 51, 52). A bioactive compound from a marine actinomycete isolate, *S. kaviengensis* showed strong antiviral activity against equine encephalitis virus (53). Padilla *et al*. (54) reported the antiviral activity of the extracts from termite associated actinobacteria against bovine viral diarrhoea virus. Lee *et al*. (11) isolated an antiviral compound namely benzatatin C from *S. nitrosporeus* and reported antiviral activity against herpes simplex virus type 1 and type 2, and vesicular stomatitis virus, and that this antiviral activity was found to be mediated due to the chlorine moiety in its molecular structure. Strand *et al*. (55) isolated a bioactive compound from marine *Streptomyces* sp. with antiviral activity against adenovirus. Hasobe *et al*. (56) reported that guanine-7-N-oxide isolated from *Streptococcus* sp. was found to inhibit *in vitro* replication of fish herpes virus, rhabdovirus, and infectious pancreatic necrovirus. Furan 2-yl acetate produced by a marine *Streptomyces* spp. VITSDK1 was found to be efficient to control fish nodavirus in seabass cell line (57). All the above-mentioned studies carried out by various workers indicate that actinomycetes isolates are a potential source of antiviral compounds. In the light of these studies was carried out to screen actinomycetes isolates collected from different habitats against WSSV.

Totally 64 actinomycetes isolates collected from different environments were screened for antiviral activity against WSSV since this virus is a serious viral pathogen responsible for severe mortality and economic loss in shrimp culture industry not only in India but also in all shrimp growing countries. One among 64 isolates the isolate designated as CAHSH-2 was found to be very effective in inactivating the WSSV at the low concentration of 0.2 mg per shrimp. As observed in the present study, actinomycetes isolates were found to reduce WSSV infection in shrimp (15, 49, 50). All studies related to antiviral activity of actinomycetes isolates against WSSV currently are at a preliminary level only. A detailed study as carried out in the present study has so far not been carried out.

The ethyl acetate extract of CAHSH-2 showed strong antiviral activity against WSSV (no mortality and no signs of WSSV) at the concentration of 0.2 mg per shrimp. Different methods such as *in vitro* assay using cell lines and animal models are being followed in different laboratories to screen the natural products for antiviral activity. In the present study, the antiviral activity of ethyl acetate extract of actinomycetes isolates against WSSV was assessed in shrimp due to lack of shrimp cell lines as followed by different workers (58, 22, 23). Testing the antiviral compounds in animal models has relatively maximum predictive value and more advantages when compared to *in vitro* assay using cell lines or chicken eggs. Testing in an animal model system would help to identify both antiviral activity and antiviral agents, and also help to determine the toxicity of compounds instantaneously while screening antiviral activity. The use of an animal model for antiviral screening of natural products should have some essential features such as use of virus with minimal alteration by adaptation, use of the natural route of infection and size of inoculum, and similarity of infection, pathogenesis, host response and compound toxicity (59).

The ethyl acetate extract of CAHSH-2 was found to be highly efficient to inactivate WSSV at the concentration of 200 μg per shrimp and this concentration was found to be non-lethal to the shrimp. The potential ethyl acetate extract was subjected to silica gel column and five fractions were obtained. Among the five fractions, fraction-F1 was found to be active against WSSV and further analysis of this active fraction resulted in the isolation of two active sub-fractions. Two different phthalate esters, namely di-n-octyl phthalate and bis (2-methylheptyl) phthalate were purified from active fraction and the structural elucidation of these compounds was confirmed by FT-IR, GC-MS and NMR spectra analyses. Among these compounds, di-n-octyl phthalate was found to more efficiency to inactive WSSV at the concentration of 50 μg per shrimp and 78% survival was found in WSSV challenged shrimp at 10 d p.i. whereas 89% survival was found in WSSV-challenged shrimp at the concentration of 200 μg per shrimp. As observed in the present study, different antiviral compounds against the viruses of human, animals and plants were reported by many workers. Nakagawa *et al*. (60) isolated and identified pentalactones from *Streptomyces* sp. M-2718 and reported that they were active against several DNA viruses. The antiviral activity of guanine-7-N-oxide produced by *Streptomyces* sp. against fish herpes virus, rhabdovirus and IPNV was tested and it was found to inhibit the replication of these viruses (56). The results obtained in the present study on potential antiviral compounds against WSSV from actinomycetes isolate agree with previous works on antiviral compounds against viruses of human, animals and fish. Hence, this potential actinomycetes isolate may be useful when incorporated in the feed to control WSSV infection in the culture system.

Two compounds, di-n-octyl phthalate and bis (2-methylheptyl) phthalate were isolated from an endophytic actinomycetes isolate and were found to be effective against WSSV of shrimp. As observed in the present study, the antiviral activity of these compounds was also reported by many workers. Elnaby *et al*. (61) tested the antiviral activity of di-n-octyl phthalate isolated from a marine actinomycetes isolate (*S. parvus*) against hepatitis C virus (HCV). Compounds produced by *Pseudoalteromonas piscida* contained benzene 1-pentyloctyl, benzene 1-butylheptyl and di-n-octyl phthalate, and these compounds were found to be active against HCV (62). Yin *et al*. (63) identified di-n-octyl phthalate in the crude extract from the endophytic fungus *Aspergillus terreus* MP15 of *Swietenia macrophylla* leaf and reported antibacterial activity against food-borne bacteria. In the present study, antiviral activity of di-n-octyl phthalate isolated from endophytic actinomycetes isolate against WSSV was reported for the first time. The anti-WSSV compound namely bis (2-methylheptyl) phthalate isolated from actinomycetes isolate in the present study was also reported to be present in the leaves of *Pongamiapinnata* and effective against WSSV (64).

The present study provides a suitable 3D model for di-n-octyl phthalate with VP26 or VP28 and bis (2-methylheptyl) phthalate with VP26 or VP28 complex formation. The ligand binding site on target proteins VP26 and VP28 were predicted using FTsite. We predicted three ligand binding sites for these target proteins. A significant variation in these three-ligand binding sites was observed (65). VP26 and VP28 are the major structural proteins of the outer membrane of WSSV of shrimp. Based on *in vivo* neutralization experiments on WSSV in *P. monodon* with antibodies raised against VP28, it is suggested that VP28 is located in the “spikes” of the WSSV envelope and this protein may thus be involved in the systemic infection of WSSV in shrimp (66, 67) and also play an important role in the process of infection in shrimp. Neutralizing the VP28 protein using anti-VP28 or silencing the genes of VP28 and VP26 using gene specific dsRNA increased the survival of WSSV-challenged shrimp (66, 67, 68, 69). These studies clearly indicate that the blocking of VP28 or VP26 would help to protect the shrimp from WSSV infection. The high survival of shrimp injected with CAHSH-2-treated WSSV might be due to the blocking of VP28 and VP26 proteins by the compounds [di-n-octyl phthalate and bis (2-methylheptyl) phthalate] present in the actinomycetes isolate (CAHSH-2) as evidenced from the docking studies. Among these two compounds, di-n-octyl phthalate was found to be more efficient to inactive WSSV in *in vivo* experiments when compared to bis (2-methylheptyl) phthalate. This observation is further confirmed by the docking studies which showed that di-n-octyl phthalate has high binding stability with VP26 and VP28 whereas the bis (2-methylheptyl) phthalate has low binding affinity with VP26 and VP28. High binding stability with VP28 and VP26 might be the reason for high efficiency of di-n-octyl phthalate to inactivate the WSSV when compared to bis (2-methylheptyl) phthalate. Similar observation was made by Sundharsana *et al*. (70) in *in silico* molecular docking and simulation analysis of VP26 and VP28 proteins of WSSV with the ligand 3-(1-chloropiperidin-4-yl)-6-fluoro benzisoxazole 2 and the results revealed the ligand binding with polar amino acids in the pore region of viral proteins.

In the present study, the antiviral activity of potential actinomycetes isolate, active fractions and pure antiviral compounds against WSSV was confirmed by various methods such as bioassay test, PCR, RT-PCR, Western blot, indirect ELISA and real time PCR. In the previous works, the antiviral activity of actinomycetes isolates against WSSV was confirmed by either bioassay test, PCR assay or both (15, 49, 50). The results of all confirmatory assays indicate the potentiality of actinomycetes isolates to inactivate the WSSV as observed by Kumar *et al*. (15) and Jenifer *et al*. (49).

Before the application of antiviral drugs from actinomycetes or any other organism, they should be tested for toxicity to prove that it is non-toxic to the host and the environment (71, 72). In the present study, the toxicity of ethyl acetate extract of actinomycetes isolate (CAHSH-2) and antiviral compound (di-n-octyl phthalate) was tested *in vivo* using *Artemia* nauplii, post-larvae and adult shrimp, and *in vitro* assay using fish cell lines. The results of *in vivo* assays revealed that high doses of extract or active compound did not produce mortality in *Artemia*, post-larvae and adult shrimp, and this indicates that the extract of potential actinomycetes isolate and active compound are safe for use in shrimp culture system as observed by other workers (73, 74, 74). Das *et al*. (73) reported that *Streptomyces* sp. is not toxic to both nauplii and adults of *A. salina*. More studies revealed that *Streptomyces* strains from different environments were harmless and did not result in any sign of abnormal behavior or mortality in *P. monodon* (74) and *L. vannamei* (75).

Various characters like morphological, physiological and biochemical characters are used in the classical approach to identify the actinomycetes isolates. The classical method of classification described in the identification key by Nonomura (77) and Bergey’s Manual of Determinative Bacteriology (78) was found to be useful to classify the actinomycetes isolates in the present study. All the characteristics studied for CAHSH-2 in the present study were compared with that of reference strain, *Streptomyces ghanaensis* obtained from MTCC, Chandigarh, India, and showed 94% similarity. In addition to these characteristics, molecular tools were applied to bacterial taxonomy like other organisms (78, 79). In the present study, the 16S RNA method was followed to identify the CAHSH-2 isolate. Nucleotide sequences for the 16S ribosomal RNA (rRNA) genes are available for almost all actinomycete species and reference to these sequences allow for easy identification of actinomycetes isolates. The 16S RNA gene sequence of CAHSH-2 isolate exhibited sequence similarity of 99% with *S. ghanaensis*. Based on morphological, physiological, biochemical and molecular taxonomy, the CAHSH-2 isolate was identified as *Streptomyces ghanaensis-like* strain and deposited in the repository of C. Abdul Hakeem College with the designation *Streptomyces ghanaensis-CHASH-2* for scientific purpose.

## Acknowledgement

The authors are grateful to the Management of C. Abdul Hakeem College, Melvisharam, India, for providing the facilities to carry out this work. This study was funded by the Department of Biotechnology (**BT/PR4132/AAQ/3/576/2011**), Government of India, New Delhi, India.

